# Complex genetic admixture histories reconstructed with Approximate Bayesian Computations

**DOI:** 10.1101/761452

**Authors:** Cesar A. Fortes-Lima, Romain Laurent, Valentin Thouzeau, Bruno Toupance, Paul Verdu

## Abstract

Admixture is a fundamental evolutionary process that has influenced genetic patterns in numerous species. Maximum-likelihood approaches based on allele frequencies and linkage-disequilibrium have been extensively used to infer admixture processes from dense genome-wide datasets mostly in human populations. Nevertheless, complex admixture histories, beyond one or two pulses of admixture, remain methodologically challenging to reconstruct, especially when large datasets are unavailable. We develop an Approximate Bayesian Computations (ABC) framework to reconstruct complex admixture histories from independent genetic markers. We built the software package *MetHis* to simulate independent SNPs in a two-way admixed population for scenarios with multiple admixture pulses, or monotonically decreasing or increasing admixture at each generation; drawing model-parameter values from prior distributions set by the user. For each simulated dataset, we calculate 24 summary statistics describing genetic diversity and moments of individual admixture fraction. We coupled *MetHis* with existing ABC algorithms and investigate the admixture history of an African American and a Barbadian population. Results show that Random-Forest ABC scenario-choice, followed by Neural-Network ABC posterior parameter estimation, can distinguish most complex admixture scenarios and provide accurate model-parameter estimations. For both admixed populations, we find that monotonically decreasing contributions over time, from the European and African sources, explain the observed data more accurately than multiple admixture pulses. Furthermore, we find contrasted trajectories of introgression decay from the European and African sources between the two admixed populations. This approach will allow for reconstructing detailed admixture histories in numerous populations and species, particularly when maximum-likelihood methods are intractable.

## INTRODUCTION

Hybridization between species and admixture between populations are powerful mechanisms influencing biological evolution. Genetic admixture patterns have thus been extensively studied to understand migrations and admixture-related adaptation (HELICONIUS GENOME CONSORTIUM 2012; HELLENTHAL *et al*. 2014; SKOGLUND *et al*. 2015; BRANDENBURG *et al*. 2017). The increasing availability of genome-wide data in numerous species, and particularly humans (e.g. 1000 GENOMES PROJECT CONSORTIUM 2015), further provides unprecedented opportunities to understand the genomic architecture of admixture, characterize the contribution of admixture to adaptive evolution, and infer demographic histories of admixture from genetic data.

Based on a long history of statistical developments aimed at investigating admixture patterns from genetic data (BERNSTEIN 1931; CAVALLI-SFORZA and BODMER 1971; CHAKRABORTY and WEISS 1988; LONG 1991; FALUSH *et al*. 2003; PATTERSON *et al*. 2012), population geneticists recently developed methods to reconstruct the genomic architecture of admixed segments deriving from each source population, and to describe admixture linkage-disequilibrium (LD) patterns (SANKARARAMAN *et al*. 2008; PRICE *et al*. 2009; LAWSON *et al*. 2012; MAPLES *et al*. 2013; GUAN 2014; SALTER-TOWNSHEND and MYERS 2019). In *Homo sapiens*, these methods have been extensively used to infer populations’ ancestral genetic origins and map local ancestry along individual genomes, often for disease-mapping purposes (e.g. SHRINER *et al*. 2011). Furthermore, by coupling admixture mapping approaches with natural selection scans, sometimes accounting for ancient and recent demographic history, it is possible to identify signatures of adaptive introgression or post-admixture selection having influenced genomic diversity patterns in human populations (JEONG *et al*. 2014; RACIMO *et al*. 2015; PATIN *et al*. 2017).

In this context, several maximum-likelihood approaches have been developed to estimate the parameters of admixture models (time of admixture events and their associated intensities) that vastly improved our understanding of detailed admixture histories in particular for human populations (e.g. PICKRELL and PRITCHARD 2012; HELLENTHAL *et al*. 2014). The two classes of methods most extensively deployed in the past rely, respectively, on the moments of allelic frequency spectrum divergences among populations (REICH *et al*. 2009; PATTERSON *et al*. 2012; PICKRELL and PRITCHARD 2012; LIPSON *et al*. 2013), and on admixture LD patterns (POOL and NIELSEN 2009; MOORJANI *et al*. 2011; GRAVEL 2012; LOH *et al*. 2013; HELLENTHAL *et al*. 2014; CHIMUSA *et al*. 2018). They allow for identifying admixture events in a given set of populations, estimating admixture fractions, and inferring time since each pulse of admixture. Notably, Gravel (GRAVEL 2012) developed an approach to fit the observed curves of admixture LD decay to those theoretically expected under admixture models involving one or two possible pulses of admixture from multiple source populations. This major advance significantly improved our ability to reconstruct detailed admixture histories using genetic data, for instance among several populations descending from the Transatlantic Slave Trade (TAST) across the Americas (e.g. MORENO-ESTRADA *et al*. 2013; BAHARIAN *et al*. 2016; FORTES-LIMA *et al*. 2017).

Despite the unquestionable importance of these previous developments, existing admixture history inference methods somewhat suffer from inherent limitations acknowledged by the authors (GRAVEL 2012; LIPSON *et al*. 2013; HELLENTHAL *et al*. 2014). First, most likelihood approaches can only consider one or two pulses of admixture in the history of the hybrid population. Nevertheless, admixture processes in numerous species are known to be often much more complex, involving multiple admixture-pulses or periods of recurring admixture over time from each source population separately. It is not yet clear how these methods might behave when they can consider only simplified versions of the true admixture history underlying the observed data (GRAVEL 2012; LIPSON *et al*. 2013; LOH *et al*. 2013; HELLENTHAL *et al*. 2014; MEDINA *et al*. 2018; NI *et al*. 2019). Second, while it is possible to compare maximum-likelihood values obtained from fitting one or two admixture pulses to the observed data as a guideline to find the “best” scenario, formal statistical comparison of model posterior probabilities is often out of reach of these approaches (GRAVEL 2012; FOLL *et al*. 2015; NI *et al*. 2019). Finally, admixture-LD methods, in particular, rely on fine mapping of local ancestry segments in individual genomes and thus require substantial amounts of genomic data (typically several hundred thousand to several millions of SNPs), and, sometimes, accurate phasing. These still represent major challenges for most species, including humans.

To overcome these limitations, Approximate Bayesian Computation (ABC) approaches (TAVARÉ *et al*. 1997; PRITCHARD *et al*. 1999; BEAUMONT *et al*. 2002) represent a promising class of methods to infer complex admixture histories from observed genetic data. Indeed, ABC has been successfully used previously in different species (including humans), and using different types of genetic data, to formally test alternative demographic scenarios hypothesized to be underlying observed genetic patterns, and to estimate, a posteriori, the parameters of the winning models (VERDU *et al*. 2009; BOITARD *et al*. 2016; FRAIMOUT *et al*. 2017).

ABC model-choice and posterior parameter inference rely on comparing observed summary statistics to the same set of statistics, calculated from a usually large number of genetic simulations explicitly parametrized by the user, and produced under competing demographic scenarios (BEAUMONT *et al*. 2002; WEGMANN *et al*. 2009; BLUM and FRANÇOIS 2010; CSILLÉRY *et al*. 2012; PUDLO *et al*. 2016; SISSON *et al*. 2018). Each simulation, and corresponding vector of summary statistics, is produced using model-parameters drawn randomly from prior distributions informed adequately by the user. Therefore, the flexibility of ABC relies mostly on explicit genetic data simulations set by the user. This makes ABC *a priori* particularly well suited to investigate highly complex historical admixture scenarios for which likelihood functions are very often intractable, but for which simulation of genetic data is feasible (PRITCHARD *et al*. 1999; VERDU and ROSENBERG 2011; GRAVEL 2012). However, ABC has until now seldom been used to investigate admixture processes beyond a single admixture pulse or constant migrations (BUZBAS and ROSENBERG 2015; BUZBAS and VERDU 2018).

In this paper, we show how ABC can be successfully applied to reconstruct, from genetic data, highly complex admixture histories beyond exploring models with a single or two pulses of admixture. In particular, we focus on evaluating how a relatively limited number of independent SNPs can be used for accurately distinguishing major classes of historical admixture models, such as multiple admixture-pulses versus recurring increasing or decreasing admixture over time, and for conservative posterior parameter inference under the winning model. Furthermore, we show that the quantiles and higher moments of the distribution of admixture fractions in the admixed population are highly informative summary-statistics for ABC model-choice and posterior-parameter estimation, as expected analytically (VERDU and ROSENBERG 2011; GRAVEL 2012; BUZBAS and VERDU 2018).

In order to do so, and since genetic data simulation under highly complex admixture models is not trivial using existing coalescent approaches (WAKELEY *et al*. 2012), we propose a novel *ad hoc* forward-in-time genetic data simulator and a set of parameter-generator and summary-statistics calculation tools embedded in an open source C software package called *MetHis*. It is adapted to conduct primarily ABC inferences with existing ABC tools implemented in the R (R DEVELOPMENT CORE TEAM 2017) packages *abc* (CSILLÉRY *et al*. 2012) and *abcrf* (PUDLO *et al*. 2016; RAYNAL *et al*. 2019).

We exemplify our approach by reconstructing the complex admixture histories underlying observed genetic patterns separately for the African American (ASW) and Barbadian (ACB) populations from the 1000 Genomes Project Phase 3 (1000 GENOMES PROJECT CONSORTIUM 2015). Both populations are known to be admixed populations of European and African descent in the context of the TAST (e.g. GRAVEL 2012; BAHARIAN *et al*. 2016; MARTIN *et al*. 2017). We find admixture histories much more complex than previously inferred for these populations and further reveal the diversity of admixture histories undergone by populations descending from the TAST in the Americas.

## MATERIAL AND METHODS

We aimed at evaluating how ABC model-choice and posterior parameter estimation could allow reconstructing highly complex historical admixture processes using independent genome-wide SNPs. To do so, we chose to focus on the recent admixture history of populations of African and European ancestry, descending from European colonization and the TAST in the Americas. This case-study represents an appropriate setting for empirically testing our ABC approach, since this period of history starting in the late 15th century has been extensively studied in population genetics based on the same publicly available datasets.

First, we describe the targeted case-study population and genetic datasets. Second, we present in detail the complex admixture processes here investigated and the associated demographic parameters. Third, we describe the novel simulation and summary statistics calculation software package called *MetHis*, here proposed to investigate these admixture processes. Fourth, we detail the Random-Forest ABC procedure used for scenario-choice inference and the performance of this approach both in general for the tested models and specifically for the real data here investigated. Finally, we detail the Neural Network ABC procedure deployed to estimate posterior parameter distributions, its parameterization, and the cross-validation procedures conducted to evaluate its power and accuracy.

### Population Genetics Dataset

We considered the admixture histories of the African American (ASW) and Barbadian (ACB) population samples from the 1000 Genomes Project Phase 3 (1000 GENOMES PROJECT CONSORTIUM 2015). Previous studies identified, within the same database, the West European Great-Britain (GBR) and the West African Yoruba (YRI) population samples as reasonable proxies for the genetic sources of the admixture of both ACB and ASW populations, consistently with the macro-history of the TAST in the former British colonial empire in Africa and the Americas (BAHARIAN *et al*. 2016; MARTIN *et al*. 2017; VERDU *et al*. 2017).

We excluded from our sample set, individuals previously identified to be more closely related than first-degree cousins in the four populations separately (VERDU *et al*. 2017). We also excluded the three ASW individuals showing traces of Native American or East-Asian admixture beyond that from Europe and Africa, as reported in previous studies (MARTIN *et al*. 2017). This allows us to consider only two source populations for the admixture history of both admixed populations investigated here. Among the remaining individuals we randomly drew 50 individuals in the targeted admixed ACB and ASW populations, respectively, and included the remaining 90 YRI individuals and 89 GBR individuals.

We extracted biallelic polymorphic sites (SNPs as defined by the 1000 Genomes Project Phase 3) from the merged ACB+ASW+GBR+YRI data set, excluding singletons. Furthermore, we focused only on independent SNPs by LD pruning the data set using the PLINK (PURCELL *et al*. 2007) --indep-pairwise option with a sliding window of 100 SNPs, moving in increments of 10 SNPs, and r^2^ threshold of 0.1 (ALEXANDER *et al*. 2009). Finally, we randomly drew 100,000 SNPs from the remaining SNP set.

### Competing complex admixture scenarios

We aimed at investigating comprehensive admixture histories with, after the original foundation of the admixed population, possibly multiple pulses (>1) of admixture, or recurring monotonically increasing or decreasing admixture, from each source population separately. To do so, we chose to work under the general mechanistic model presented in Verdu and Rosenberg (VERDU and ROSENBERG 2011), henceforth called the VR2011 model, derived from Ewens and Spielman (EWENS and SPIELMAN 1995). Briefly (**Supplementary Figure S1**), the VR2011 general model considers, for diploid organisms, a panmictic admixture process, discrete in generations, where *M* source populations S_m_ contribute to the hybrid population H at the following generation *g* + 1 with proportions *S_m,g_* each in [0,1], and where the hybrid population H contributes to itself with proportion *h_g_* in [0,1] with *h*_0_ = 0, satisfying, for each value of *g* ≥ 0, ∑*_m_*_∈[1,*M*]_ *S_m_*_,*g*_ + *h_g_* = 1.

Here, we adapted the two source-populations version of the general VR2011 (*M* = 2), and define, next, the nine competing complex admixture scenarios considered to reconstruct the history of introgression from Africa and Europe into the gene-pool of the ACB and ASW admixed populations (see above), separately (**Figure 1**).

**Figure 1.**
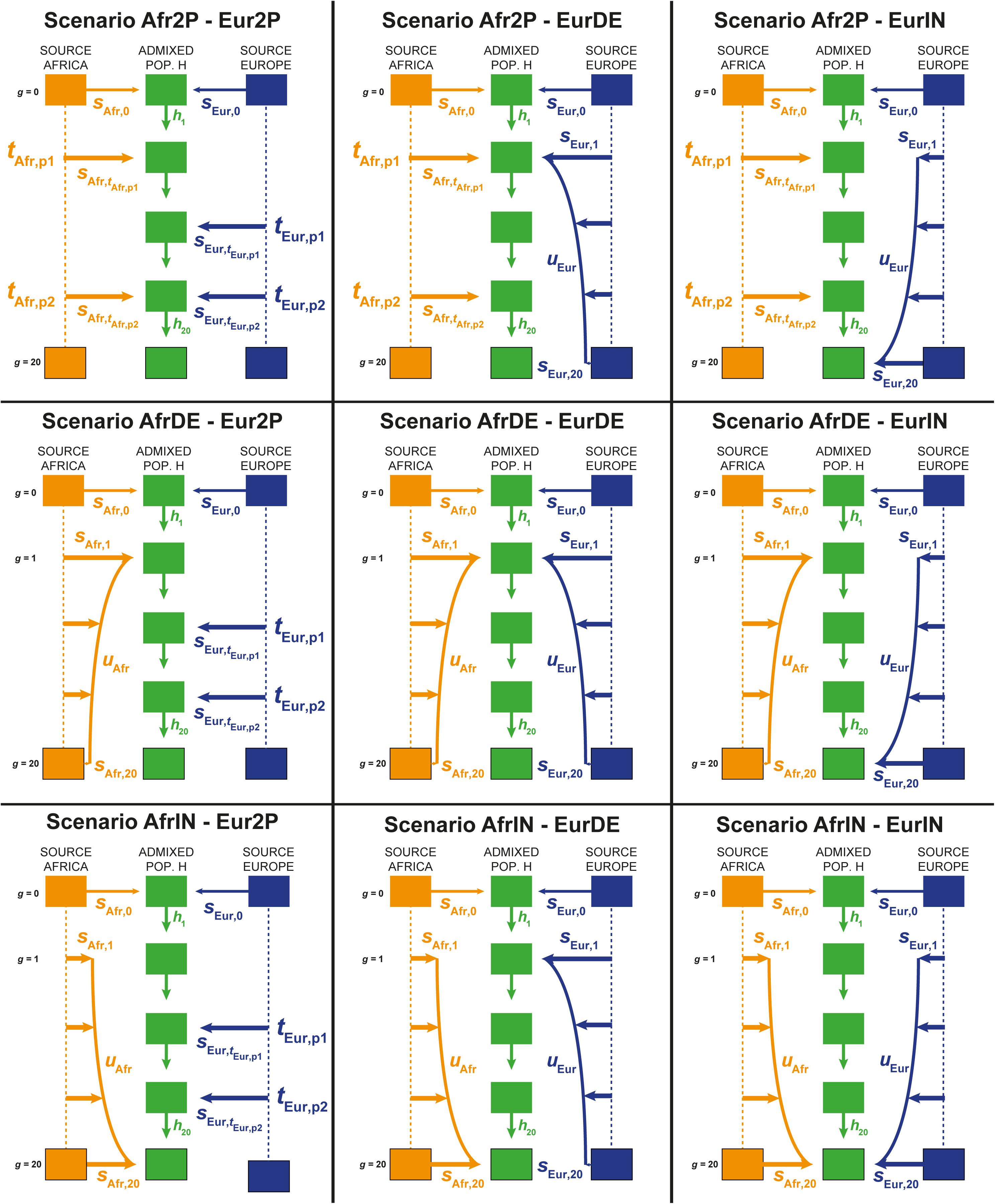
Nine competing scenarios for reconstructing the admixture history of African American ASW or Barbadian ACB populations descending from West European and West sub-Saharan African source populations during the Transatlantic Slave Trade. “EUR” represents the Western European and “AFR” represents the West Sub-Saharan African source populations for the admixed population H. See **Table 1** and **Material and Methods** for model parameter descriptions.

**Table 1.** Parameter prior distributions for simulation with *MetHis* and Approximate Bayesian Computations historical inference. Parameter list correspond to the nine competing historical admixture models described in **Figure 1** and **Material and Methods**.

| Parameter Names | Prior distribution | Condition | Models |
| --- | --- | --- | --- |
| $s_{Afr,0}$ | *Uniform* [0,1] | - | all models |
| $t_{Afr,p1}$<br>$t_{Afr,p2}$ | *Uniform* [0,20] | $t_{Afr,p1} \neq t_{Afr,p2}$ | Afr2P models |
| $s_{Afr}, t_{Afr,p1}$<br>$s_{Afr}, t_{Afr,p2}$ | *Uniform* [0,1] | For all $g$, $h_g = 1 - s_{Afr,g} - s_{Eur,g}$ in [0,1] | Afr2P models |
| $t_{Eur,p1}$<br>$t_{Eur,p2}$ | *Uniform* [0,20] | $t_{Eur,p1} \neq t_{Eur,p2}$ | Eur2P models |
| $s_{Eur}, t_{Eur,p1}$<br>$s_{Eur}, t_{Eur,p2}$ | *Uniform* [0,1] | For all $g$, $h_g = 1 - s_{Afr,g} - s_{Eur,g}$ in [0,1] | Eur2P models |
| $s_{Afr,1}$ | *Uniform* [0,1] | For all $g$, $h_g = 1 - s_{Afr,g} - s_{Eur,g}$ in [0,1] | AfrDE models |
| $s_{Afr,20}$ | *Uniform* [0, $s_{Afr,1} / 3$] | For all $g$, $h_g = 1 - s_{Afr,g} - s_{Eur,g}$ in [0,1] | AfrDE models |
| $u_{Afr}$ | *Uniform* [0,0.5] | - | AfrDE models |
| $s_{Eur,1}$ | *Uniform* [0,1] | For all $g$, $h_g = 1 - s_{Afr,g} - s_{Eur,g}$ in [0,1] | EurDE models |
| $s_{Eur,20}$ | *Uniform* [0, $s_{Eur,1} / 3$] | For all $g$, $h_g = 1 - s_{Afr,g} - s_{Eur,g}$ in [0,1] | EurDE models |
| $u_{Eur}$ | *Uniform* [0,0.5] | - | EurDE models |
| $s_{Afr,1}$ | *Uniform* [0, $s_{Afr,20} / 3$] | For all $g$, $h_g = 1 - s_{Afr,g} - s_{Eur,g}$ in [0,1] | AfrIN models |
| $s_{Afr,20}$ | *Uniform* [0,1] | For all $g$, $h_g = 1 - s_{Afr,g} - s_{Eur,g}$ in [0,1] | AfrIN models |
| $u_{Afr}$ | *Uniform* [0,0.5] | - | AfrIN models |
| $s_{Eur,1}$ | *Uniform* [0, $s_{Eur,20} / 3$] | For all $g$, $h_g = 1 - s_{Afr,g} - s_{Eur,g}$ in [0,1] | EurIN models |
| $s_{Eur,20}$ | *Uniform* [0,1] | For all $g$, $h_g = 1 - s_{Afr,g} - s_{Eur,g}$ in [0,1] | EurIN models |
| $u_{Eur}$ | *Uniform* [0,0.5] | - | EurIN models |

#### Foundation of the admixed population H

For all scenarios (**Figure 1**, **Table 1**) we chose a fixed time for the foundation (generation 0, forward-in-time) of population H occurring 21 generations before present, with admixture proportions *s*_Afr,0_ and *s*_Eur,0_ from the African and the European sources respectively, with *s*_Afr,0_ + *s*_Eur,0_ = 1, and *s*_Afr,0_ in [0,1]. This corresponds to the first arrival of European permanent settlers in the Americas and Caribbean in the late 15^th^ and early 16^th^ centuries, considering 20 or 25 years per generation and the sampled generation born in the 1980s. Note that simulations considering a parameter *s*_Afr,0_ close to 0, or alternatively 1, correspond to foundations of the population H from either one source population, therefore delaying the first “real” genetic admixture event to the next, more recent, demographic event. Following foundation, we consider three alternative scenarios for the admixture contribution of each source population S, African or European in our case, separately.

#### Admixture-pulse(s) scenarios

For a given source population S, African (Afr) or European (Eur), scenarios *S-2P* consider two possible pulses of admixture into population H occurring respectively at time *t*_S,p1_ and *t*_S,p2_ distributed in [1,20] with *t*_S,p1_ ≠ *t*_S,p2_, with associated admixture proportion *s*_S,tS,p1_ and *s*_S,tS,p2_ in [0,1] satisfying, at all times *t*, ∑*_s_*_∈(*Afr*,*Eur*)_ *s_s,t_*≤1 (**Figure 1**, **Table 1**). Note that for one of either *s*_S,t_ parameter values close to 0, the two-pulse scenarios are equivalent to single pulse scenarios after the foundation of H. Furthermore, for both *s*_S,t_ values close to 0, scenarios *S-2P* are nested with scenarios where only the founding admixture pulse 21 generations ago is the source of genetic admixture in population H. Alternatively, *s*_S,t_ parameter values close to 1 consider a virtual complete genetic replacement of population H by source population S at that time. Finally, certain *S-2P* scenarios with two consecutive pulses from a given source S (*t*_S,p1_ = *t*_S,p2_ - 1), may be strongly resembling single-pulse scenarios (after foundation).

#### Recurring decreasing admixture scenarios

For a given source population S, scenarios *S-DE* consider a recurring monotonically decreasing admixture from source population S at each generation between generation 1 (after foundation at generation 0) and generation 20 (sampled population) (**Figure 1**, **Table 1**). In these scenario, *s*_S,g_, with *g* in [1..20], are the discrete numerical solutions of a rectangular hyperbola function over the 20 generations of the admixture process until present as described in **Supplementary Note S1**. In brief, this function is determined by parameter *u*_S_, the “steepness” of the curvature of the decrease, in [0,1/2], *s*_S,1_, the admixture proportion from source population S at generation 1 (after foundation), in [0,1], and *s*_S,20_, the last admixture proportion in the present, in [0,*s*_S,1_/3]. Note that we chose the boundaries for *s*_S,20_ in order to reduce the parameter space and nestedness among competing scenarios, and explicitly force scenarios *S-DE* into a substantially decreasing admixture process. Indeed, defining *s*_S,20_ in [0,*s*_S,1_] instead would have also allowed for both decreasing admixture processes and relatively constant recurring admixture processes. Furthermore, note that parameter *u*_S_ values close to 0 create pulse-like scenarios occurring immediately after foundation of intensity *s*_S,1_, followed by constant recurring admixture at each generation until present of intensity *s*_S,20_. Alternatively, parameter *u*_S_ values close to 1/2 create scenarios with a linearly decreasing admixture between *s*_S,1_ and *s*_S,20_ from source population S at each generation after the foundation of population H.

#### Recurring increasing admixture scenarios

Finally, for a given source population S, scenarios *S-IN* mirrors the *S-DE* scenarios by considering instead a recurring monotonically increasing admixture from source population S (**Figure 1**, **Table 1**). Here, *s*_S,g_, with *g* in [1..20], are the discrete numerical solutions of the same function as in the S-DE decreasing scenarios (see above), flipped over time between generation 1 and 20. In these scenarios, *s*_S,20_ is defined in [0,1] and *s*_S,1_ in [0,*s*_S,20_/3], and *u* in [0,1/2] parametrizes the “steepness” of the curvature of the increase. Note that in this case, parameter *u* values close to 0 create pulse-like scenarios occurring in the present of intensity *s*_S,20_, preceded by constant recurring admixture of intensity *s*_S,1_ at each generation since foundation. Alternatively, parameter *u_S_* values close to 1/2 create scenarios with a linearly increasing admixture between *s*_S,1_ and *s*_S,20_ from source population S at each generation after the foundation of population H.

#### Combining admixture scenarios from either source populations

We combine these three scenarios to obtain nine alternative scenarios with two source populations, African (Afr) and European (Eur) respectively, for the admixture history of population H (**Figure 1**, **Table 1**), the ASW or ACB alternatively, with the only condition that, at each generation *g* in [1..20], parameters satisfy *s*_Afr,g_ + *s*_Eur,g_ + *h*_g_ = 1, with *h*_g_, in [0,1], being the remaining contribution of the admixed population H to itself at the generation *g*. Four scenarios (Afr2P-EurDE, Afr2P-EurIN, AfrDE-Eur2P, and AfrIN-Eur2P) consider a mixture of pulse-like and recurring admixture from each source. Three scenarios (Afr2P-Eur2P, AfrDE-EurDE, and AfrIN-EurIN), consider symmetrical classes of admixture scenarios from either source. Two scenarios (AfrIN-EurDE and AfrDE-EurIN) consider mirroring recurring admixture processes. Importantly, this scenario design considers nested historical scenarios in specific parts of the parameter space, as exemplified above.

### Forward-in-time simulations with *MetHis*

Simulation of genome-wide independent SNPs under highly complex admixture histories is often not trivial under the coalescent and using classical existing software (WAKELEY *et al*. 2012). In this context, we developed *MetHis*, a C open-source software package available at https://github.com/romain-laurent/MetHis, to simulate large amounts of genetic data under the two-source populations VR2011 model and calculate summary statistics of interest to population geneticists interested in complex admixture processes. *MetHis*, in its current form, can be used to simulate any number of independent SNPs in the admixed population H. However, *MetHis* does not allow simulating the source populations for the admixture process. Instead, this can be done efficiently using coalescent-based simulations with existing software such as *fastsimcoal2* (EXCOFFIER and FOLL 2011; EXCOFFIER *et al*. 2013), or other forward-in-time genetic data simulators such as *SLIM v*3 (HALLER and MESSER 2019).

#### Simulating source populations

Here, we wanted to focus our investigation specifically on the admixture process undergone by the admixed population descending from the TAST. Therefore, we made several *ad hoc* simplification choices for simulating source population genetic data under the nine competing models described next.

We consider that the African and European populations at the source of the admixture processes are very large populations at the drift-mutation equilibrium, accurately represented by the Yoruban YRI and British GBR datasets here investigated. Therefore, we first build two separate datasets each comprising 20,000 haploid genomes of 100,000 independent SNPs, each SNP being randomly drawn in the site frequency spectrum (SFS) observed for the YRI and GBR datasets respectively. These two datasets are used as fixed gamete reservoirs for the African and European source population datasets separately, at each generation of the forward-in-time admixture process. From these reservoirs, at each generation separately, we build an effective individual gene-pool of diploid size *N*_g_ (see below), by randomly pairing gametes avoiding selfing. These virtual source populations provide the parental pool for simulating individuals in the admixed population H, at each generation separately. Thus, while our gamete reservoirs are fixed over the 21 generations of the admixture processes here considered, the parental genetic pools are randomly built anew at each generation of the admixture process.

#### Simulating the admixed population

At each generation, *MetHis* performs simple Wright-Fisher (FISHER 1922; WRIGHT 1931) forward-in-time simulations, individual-centered, in a panmictic admixed population H of diploid effective size *N_g_*. For a given individual in the hybrid population at the following generation (*g* + 1), *MetHis* independently draws each parent from the source populations with probability *S_S,g_* (**Figure 1**, **Table 1**), or from the hybrid population with probability *h_g_*, randomly builds a haploid gamete of 100,000 independent SNPs for each parent, and pairs the two constructed gametes to create the new individual. Here, we decided to neglect mutation over the 21 generations of admixture considered. This is reasonable when studying relatively recent admixture histories. Nevertheless, this will be improved in future versions of the software, in particular to allow studying much more ancient admixture histories. Finally, while we chose explicitly to simulate only the individuals in the admixed population H here, note that future developments of *MetHis* will allow to also simulate individual genetic data in the source populations in the same way.

#### Effective population size in the source and the admixed populations

To focus on the admixture process itself without excessively increasing the parameter space, we consider, for each nine-competing model, both source populations and the admixed population H with constant effective population size *N_g_* = 1000 diploid individuals at each generation. Nevertheless, note that *MetHis* software readily allows the user to easily parameterize changes in the effective size of population H at each generation.

#### Sampling simulated unrelated individuals

After each simulation, we randomly draw individual samples matching sample sizes in our observed dataset: 90 and 89 individuals respectively from the African and European sources, and 50 individuals in the admixed population H. We sample individuals until our sample set contains no individuals related at the 1^st^ degree cousin within each population and between the admixed population and either source populations, based on explicit parental flagging during the last 2 generations of the simulations.

#### Simulating by randomly drawing parameter values from prior distributions

With this implementation of *MetHis*, we performed 10,000 independent simulations under each nine competing scenarios described above and in **Figure 1**, drawing the corresponding model-parameters (pulse-times and associated admixture intensities, “steepness” of the recurring admixture-increases or decreases and associated initial and final admixture intensities), in prior-distributions detailed in **Table 1**. Although the user can perform *MetHis* simulations with an external parameter list, we readily provide *ad hoc* scripts in *MetHis*, which allow to easily generate parameter lists for a large number of complex admixture scenarios set by the user.

For the best models identified using Random-Forest ABC model-choice approach (PUDLO *et al*. 2016) for the ACB and ASW admixed populations respectively (see **Results**), we conducted an additional 90,000 independent simulations with the same parameter priors as in the 10,000 simulations already conducted. Thus, we considered 100,000 simulations for the best scenarios for the ACB and ASW respectively, to be used for ABC posterior parameter inference (see below).

### Summary Statistics

We considered 24 summary statistics for ABC model-choice and posterior parameter inference, computed on each simulated dataset with *MetHis*. Four statistics were strictly within-populations; four statistics were strictly between-populations; and 16 statistics were specifically calculated to describe the distribution of admixture among individuals within the admixed population H. Indeed, previous theoretical works have shown that this distribution and all its moments carried signatures of the underlying complex historical process (VERDU and ROSENBERG 2011; GRAVEL 2012). Numerous descriptive statistical approaches have been successfully developed to estimate admixture fractions from genetic data in admixed populations (e.g. ALEXANDER *et al*. 2009; PATTERSON *et al*. 2012; PICKRELL and PRITCHARD 2012). However, most methods remain computationally costly when iterated for large to very large sets of simulated genetic data. Therefore, only a few previous ABC historical inference approaches have considered the distribution of admixture fraction as a summary statistics (BUZBAS and ROSENBERG 2015; BUZBAS and VERDU 2018), although some admixture-related statistics have been embedded in ABC software packages (CORNUET *et al*. 2014).

#### Distribution of admixture fractions as a summary statistic

We estimated individual admixture distribution based on allele-sharing-dissimilarity (ASD) (BOWCOCK *et al*. 1994) and multidimensional scaling (MDS) (PASCHOU *et al*. 2007; PRICE *et al*. 2009). For each simulated dataset, we first calculated a pairwise inter-individual ASD matrix using *asd* software (https://github.com/szpiech/asd) on all pairs of sampled individuals and all 100,000 independent SNPs. Then we projected in two dimensions this pairwise ASD matrix with classical unsupervised metric MDS using the *cmdscale* function R (R DEVELOPMENT CORE TEAM 2017). We expect individuals in population H to be dispersed along an axis joining the centroids of the two proxy source populations on the two-dimensional MDS plot. We projected individuals orthogonally on this axis, and calculate individual’s relative distance to each centroid. We considered this measure to be an estimate of individual average admixture level from either source population. Note that by doing so, some individuals might show “admixture fractions” higher than one, or lower than zero, as they might be projected on the other side of the centroid when being genetically close to 100% from one source population or the other. Under an ABC framework, this is not a difficulty since this may happen also on the real data *a priori*, and our goal is to use summary statistics that mimic the observed ones. This individual admixture estimation method has been shown to be highly concordant with cluster membership fractions as estimated with ADMIXTURE (ALEXANDER *et al*. 2009) in real data analyses (e.g. VERDU *et al*. 2017). Considering the real data here investigated, we confirm these previous findings since we obtain a Spearman correlation (calculated using the *cor.test* function in *R*), of rho = 0.950 (p-value < 2.10^-16^) and rho= 0.977 (p-value < 2.10^-16^) between admixture estimates based on ASD-MDS and on ADMIXTURE, for the ACB and ASW respectively (**Supplementary Figure S2**).

We used the mean, mode, variance, skewness, kurtosis, minimum, maximum, and all 10%-quantiles of the admixture distribution obtained this way in population H, as 16 separate summary statistics for further ABC inference.

#### Within population summary statistics

We calculated SNP by SNP heterozygosities (NEI 1978) using *vcftools* (DANECEK *et al*. 2011), and considered the mean and variance of this quantity across SNPs in the admixed population as two separate summary statistics for ABC inference. Note that, these quantities are fixed for each source population, respectively, and thus uninformative in our case study, since source populations are simulated only once and used for all subsequent simulations under the nine competing models (see above).

In addition, as we computed the individual pairwise ASD matrix for calculating the distribution of admixture fraction (see above), we also considered the mean and variance of ASD values across pairs of individuals within the admixed population H, as two within-population summary statistics.

#### Between populations summary statistics

In addition to previous summary statistics, we considered multilocus pairwise *F_ST_* (WEIR and COCKERHAM 1984) between population H and each source population respectively, calculated using *vcftools* (DANECEK *et al*. 2011). Note that the *F_ST_* between the source populations is fixed, since simulated source populations are themselves fixed (see above), and thus uninformative in our case study. Furthermore, we calculated the mean ASD between individuals in population H and, separately, individuals in either source population. Finally, we computed anew from Patterson (PATTERSON *et al*. 2012) the *f_3_* statistics based on allelic frequencies obtained with *vcftools* (DANECEK *et al*. 2011). In the two-source population case, this statistic is extensively employed to test the original source of the admixture of a target admixed population, infer the time since admixture, and estimate admixture intensities using maximum-likelihood approaches.

### Approximate Bayesian Computations

*MetHis* has been designed to operate under an ABC framework for model choice and parameter inference. Thus, it allows simulating genetic data under numerous possible models by drawing parameter values in a priori distributions set by the user in a flexible way. In addition, *MetHis* allows for calculating numerous summary statistics a priori of interest to admixture processes, and provides, as outputs, scenarios-parameter vectors and corresponding summary-statistics vectors in reference tables ready to be used with the machine-learning ABC *abc* (CSILLÉRY *et al*. 2012), and *abcrf* (PUDLO *et al*. 2016; RAYNAL *et al*. 2019) *R* packages (R DEVELOPMENT CORE TEAM 2017).

#### Prior-checking

We evaluated, a priori, if the above simulation design and novel tools can simulate genetic data for which summary statistics are coherent with those observed for the ACB and ASW as the targeted admixed population. To do so, we first plotted each prior summary statistics distributions and visually verified that the observed summary statistics for the ACB and ASW respectively fell within the simulated distributions (**Supplementary Figure S3**). Second, we explored the first four PCA axes computed with the *princomp* function in *R*, based on the 24 summary statistics and all 90,000 total simulations preformed for the nine competing scenarios, and visually checked that observed summary statistics were within the cloud of simulated statistics (**Supplementary Figure S4**). Finally, we performed a goodness-of-fit approach using the *gfit* function from the *abc* package in *R*, with 1,000 replicates and tolerance level set to 0.01 (**Supplementary Figure S5**).

#### Model-choice with Random-Forest Approximate Bayesian Computation

We used Random-Forest ABC (RF-ABC) for model-choice implemented in the *abcrf* function of the *abcrf R* package to obtain the cross-validation table and associated prior error rate using an out-of-bag approach (**Figure 2**). We considered the same prior probability for the nine competing models each represented by 10,000 simulations in the reference table. For the ACB and ASW observed data separately, we performed model-choice prediction and estimation of posterior probabilities of the winning model using the *predict.abcrf* function in the same *R* package, using the complete simulated reference table for training the Random-Forest algorithm (**Figure 3**, **Supplementary Table S1**). Both sets of analyses were performed considering 1,000 decision trees in the forest after visually checking that error-rates converged appropriately (**Supplementary Figure S6**), using the *err.abcrf* function in the *R* package *abcrf*. Each summary statistics relative importance to the model-choice cross-validation was computed using the *abcrf* function (**Figure 2**). RF-ABC cross-validation procedures using groups of scenarios were conducted using the group definition option in the *abcrf* function (ESTOUP *et al*. 2018).

**Figure 2:**
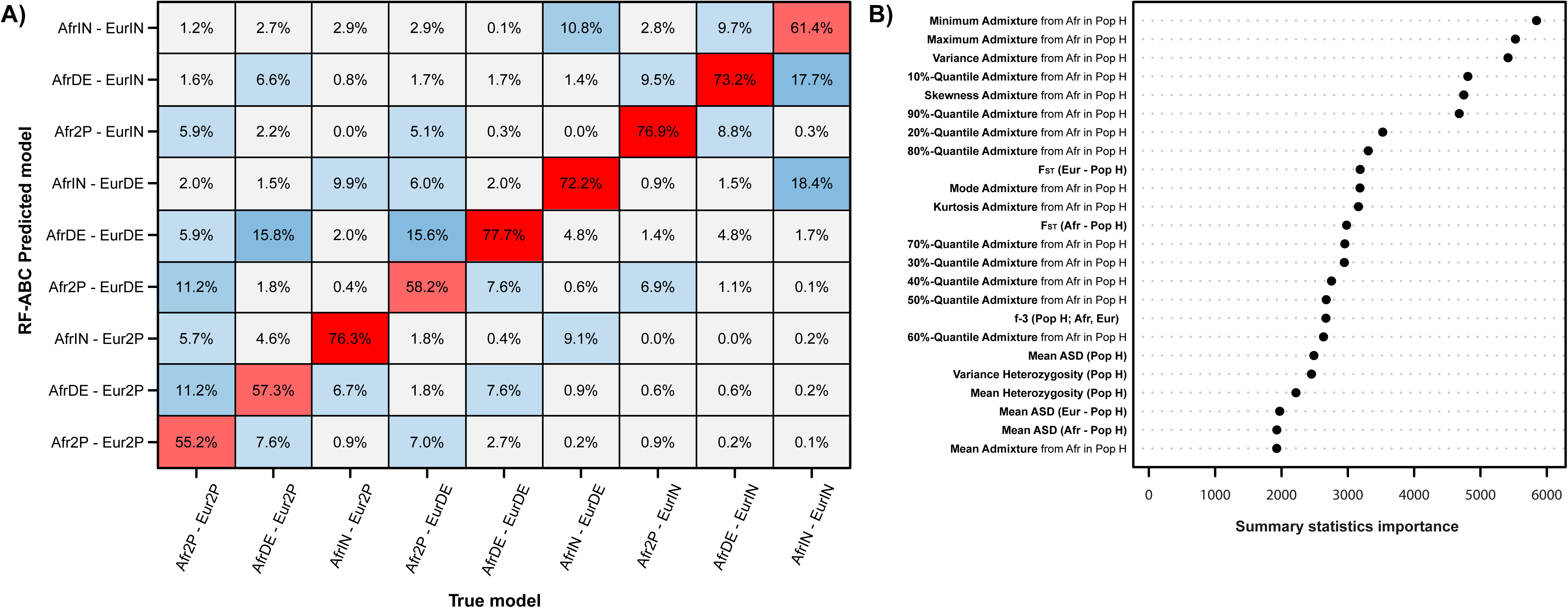
Random-Forest Approximate Bayesian Computation model-choice cross-validation. (A) Heat map of the out-of-bag cross-validation results considering each 10,000 simulations per each nine competing models (Figure 1, **Table 1**) in turn as pseudo-observed target for RF-ABC model-choice. Out-of-bag prior error rate is 32.41%. RF-ABC model-choice performed using 1,000 decision trees and 24 summary-statistics (see **Material and Methods**). (B) Summary statistics’ respective importance in the RF-ABC model-choice out-of-bag cross-validation (Pudlo et al. 2016).

**Figure 3:**
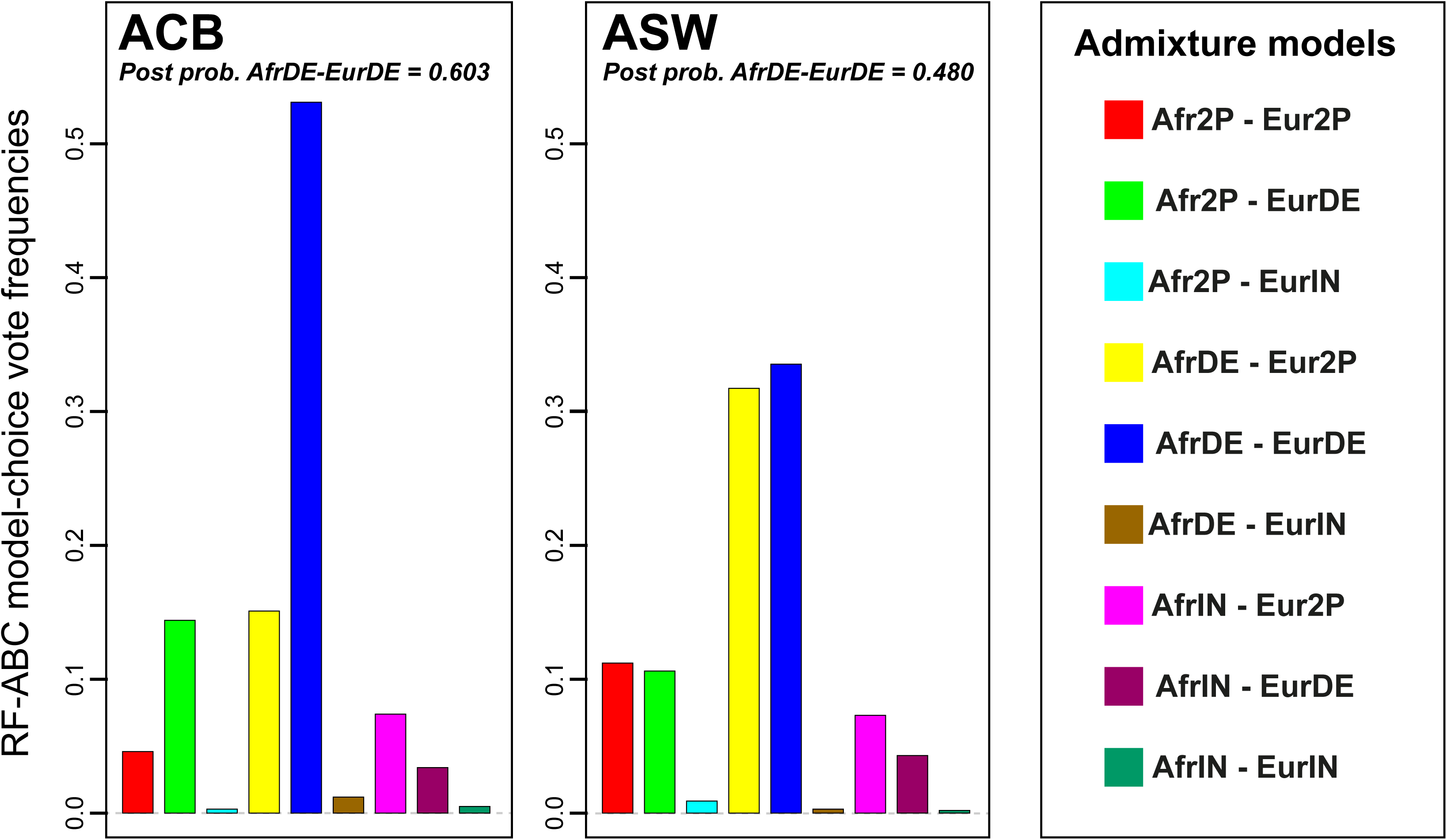
Random-Forest Approximate Bayesian Computation model-choice predictions for the ACB (left panel) and ASW (right panel) populations. Nine competing models were compared, each with 10,000 simulations (Figure 1, **Table 1**). 1,000 decision trees were considered in the model-choice prediction, respectively for each population.

#### Posterior parameter estimation with Neural-Network Approximate Bayesian Computation

It is difficult to estimate jointly the posterior distribution of all model parameters with RF-ABC (RAYNAL *et al*. 2019). Furthermore, although RF-ABC performs satisfactorily well with an overall limited number of simulations under each model (PUDLO *et al*. 2016), posterior parameter estimation with other ABC approaches, such as simple rejection (PRITCHARD *et al*. 1999), regression (BEAUMONT *et al*. 2002; BLUM and FRANÇOIS 2010) or Neural-Network (NN) (CSILLÉRY *et al*. 2012), require substantially more simulations a priori. Therefore, we performed 90,000 additional simulations, for a total of 100,000 simulations for the best scenarios identified with RF-ABC among the nine competing models for the ACB and ASW separately.

#### Neural-Network tolerance level and number of neurons in the hidden layer

For each parameter estimation analysis, we determined empirically the NN tolerance level (i.e. the number of simulations to be included in the NN training), and number of neurons in the hidden layer. Indeed, while the NN needs a substantial amount of simulations for training, there is also a risk of overfitting posterior parameter estimations when considering too large a number of neurons in the hidden layer. However, there are no absolute rules for choosing both numbers (CSILLÉRY *et al*. 2012; JAY *et al*. 2019).

Therefore, using the 100,000 simulations for the winning scenarios identified with RF-ABC (see above), we tested four different tolerance levels to train the NN (0.01, 0.05, 0.1, and 0.2), and a number of neurons ranging between four and seven (the number of free parameters in the winning scenarios, see **Results**). For each pair of tolerance level and number of neurons values, we conducted cross-validation checking of posterior parameter estimations with 1,000 randomly chosen simulated datasets in turn used as pseudo-observed data with the “*cv4abc*” function of the *R* package *abc*. We considered the median point-estimate of each posterior parameter 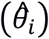 to be compared with the true parameter value used for simulation (*θ_i_*). The cross-validation parameter prediction error was then calculated across the 1,000 separate posterior estimations for pseudo-observed datasets for each pair of tolerance level and number of neurons, and for each parameter 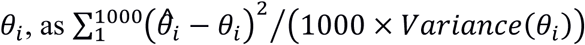 allowing to compare errors for scenarios-parameters across NN tolerance-levels and numbers of hidden neurons, using the *summary.cv4abc* function in the *R* package *abc* (CSILLÉRY *et al*. 2012). Results showed that, *a priori*, all numbers of neurons considered performed very similarly for a given tolerance level (**Supplementary Table S2**). Furthermore, results showed that considering 1% closest simulations to the pseudo-observed ones, to train the NN for parameter estimation, reduces the average error for each tested number of neurons. Thus, we decided to opt for four neurons in the hidden layer and a 1% tolerance level for training the NN in all subsequent NN-ABC analyses, in order to avoid overfitting in parameter posterior estimations.

#### Estimation of model-parameters posterior distributions for ACB and ASW

We jointly estimated model-parameters posterior distributions for the ACB and ASW admixed population separately, using 100,000 simulations for the best scenarios identified for each admixed population separately, using NN-ABC (“*neuralnet*” methods’ option in the *R* package *abc*) based on the logit-transformed (“*logit*” transformation option in the *R* package *abc*) summary statistics using a 1% tolerance level to train the NN (i.e. considering only the 1,000 closest simulations to the observed data), fitted using a single-hidden-layer neural network with four hidden neurons (**Figure 4**, **Table 2**).

**Figure 4:**
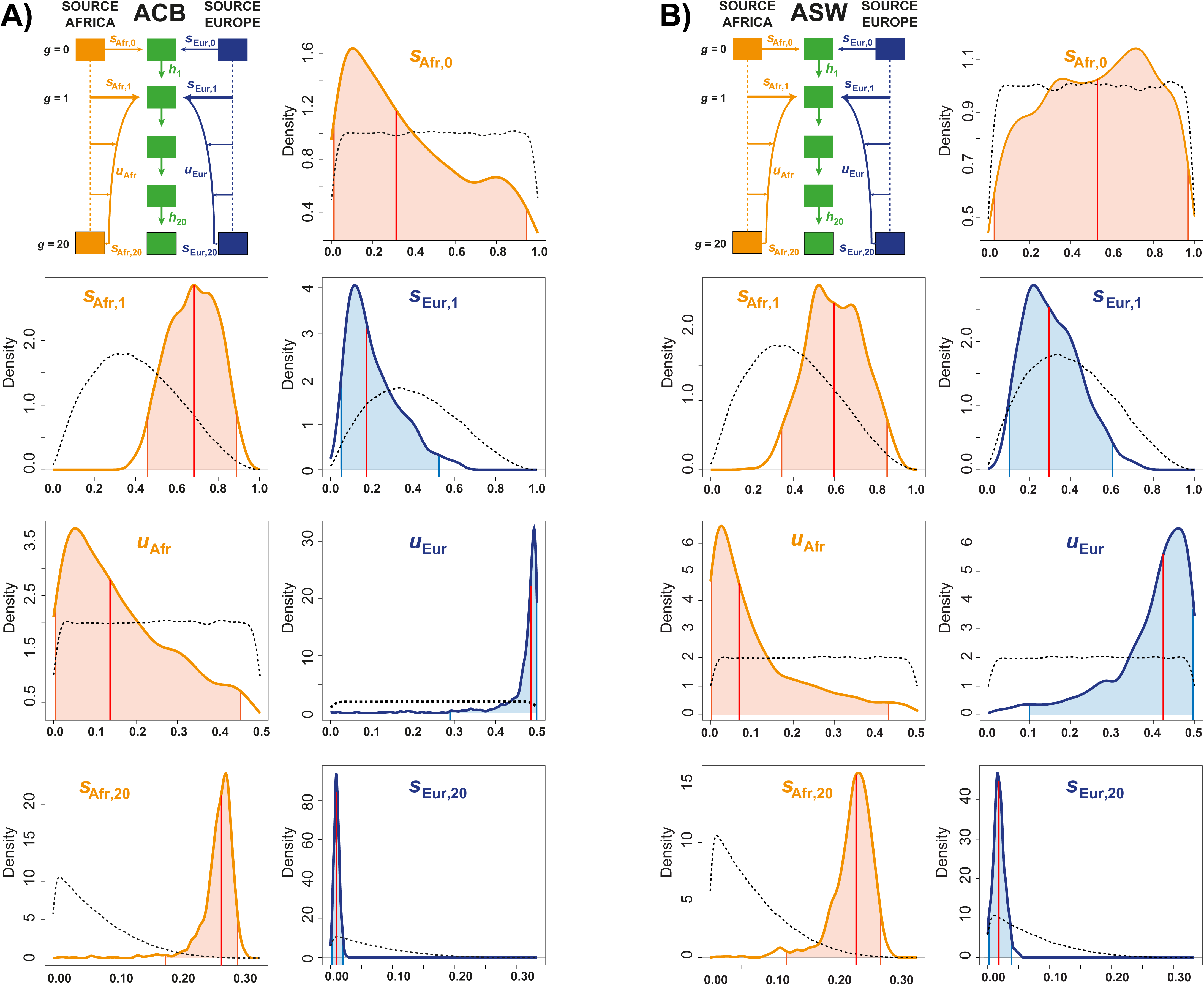
Neural-Network Approximate Bayesian Computation posterior parameters estimated densities under the winning scenario AfrDE-EurDE, for (A) the ACB and (B) the ASW populations. Median posterior point estimates are indicated by the red vertical line, 95% credibility intervals are indicated by the colored area under the posterior curve (**Table 2**). All posterior parameter estimations were conducted using 100,000 simulations under scenario AfrDE-EurDE, a 1% tolerance rate (1,000 simulations), 24 summary statistics, logit transformation of all parameters, and four neurons in the hidden layer (see **Material and Methods**). For all parameters separately, densities are plotted with 1,000 points, a Gaussian kernel, and are constrained to the prior limits. Posterior parameter densities are indicated by a solid line; prior parameter densities are indicated by black dotted lines.

**Table 2.** Neural-Network Approximate Bayesian Computation posterior parameter weighted distributions under the winning scenario AfrDE-EurDE, for the ACB and ASW populations. All posterior parameter estimations were conducted using 100,000 simulations under scenario AfrDE-EurDE (**Figure 1**, Table 1), a 1% tolerance rate (1,000 simulations), 24 summary statistics, logit transformation of all parameters, and 4 neurons in the hidden layer (see **Material and Methods**).

| *AfrDE-EurDE* |  |  |  |  |  |
| --- | --- | --- | --- | --- | --- |
|  | *parameters* | *Median* | *Mean* | *Mode* | *95% Credibility Interval* |
| **ACB** | $S_{Afr,0}$ | 0.3097 | 0.3747 | 0.1121 | [0.0116 ; 0.9347] |
| | $S_{Afr,1}$ | 0.6797 | 0.6769 | 0.6813 | [0.4577 ; 0.8880] |
| | $S_{Afr,20}$ | 0.2707 | 0.2655 | 0.2788 | [0.1985 ; 0.2967] |
| | $u_{Afr}$ | 0.1409 | 0.1684 | 0.0508 | [0.0041 ; 0.4507] |
| | $S_{Eur,1}$ | 0.1807 | 0.2160 | 0.1158 | [0.0542 ; 0.5525] |
| | $S_{Eur,20}$ | 0.0100 | 0.0102 | 0.0093 | [0.0018 ; 0.0200] |
| | $u_{Eur}$ | 0.4858 | 0.4627 | 0.4929 | [0.1886 ; 0.4992] |
| **ASW** | $S_{Afr,0}$ | 0.5258 | 0.5124 | 0.7015 | [0.0262 ; 0.9758] |
| | $S_{Afr,1}$ | 0.6006 | 0.6026 | 0.6081 | [0.3506 ; 0.8581] |
| | $S_{Afr,20}$ | 0.2352 | 0.2286 | 0.2385 | [0.1222 ; 0.2714] |
| | $u_{Afr}$ | 0.0662 | 0.1105 | 0.0253 | [0.0025 ; 0.4393] |
| | $S_{Eur,1}$ | 0.2917 | 0.3080 | 0.2203 | [0.1048 ; 0.5951] |
| | $S_{Eur,20}$ | 0.0180 | 0.0189 | 0.0157 | [0.0022 ; 0.0389] |
| | $u_{Eur}$ | 0.4250 | 0.3966 | 0.4567 | [0.1077 ; 0.4950] |

#### Posterior parameter estimation error and credibility interval accuracy

For the ACB and ASW admixed populations separately, we wanted to evaluate the posterior error performed by our NN-ABC approach on the median point estimate of each parameter, in the vicinity of our observed data rather than randomly on the entire parameter space. To do so, we first identified the 1,000 simulations closest to the real data with a tolerance level of 1%, for the ACB and ASW respectively. Then, separately for the ACB and ASW set of closest simulations, we performed, similarly as above for the real data parameter estimation procedure, 1,000 separate NN-ABC parameter estimations using the *“neural”* method in the *abc* function with a NN trained with 1% tolerance level and four neurons in the hidden layer, using in turn the other 99,999 simulations as reference table, and recorded the median point estimate for each parameter. We then compared these estimates with the true parameter used for each 1,000 pseudo-observed target in the vicinity of our observed data and provide three types of error measurements in **Table 3**. The mean-squared error scaled by the variance of the true parameter 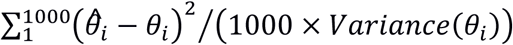 as previously (Csilléry et al. 2012); the mean-squared error 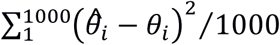, allowing to compare estimation errors for a given scenario-parameter between the ACB and ASW analyses; and the mean absolute error 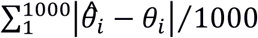, which provides a more intuitive parameter estimation error.

**Table 3.** Neural-Network Approximate Bayesian Computation posterior parameter errors under the winning scenario AfrDE-EurDE, for the ACB and ASW populations. For each target population separately, we conducted cross-validation by considering in turn 1,000 separate NN-ABC parameter inferences each using in turn one of the 1,000 closest simulations to the observed ACB (or ASW) data as the target pseudo-observed simulation. All posterior parameter estimations were conducted using 100,000 simulations under scenario AfrDE-EurDE (**Figure 1**, Table 1), a 1% tolerance rate (1,000 simulations), 24 summary statistics, logit transformation of all parameters, and four neurons in the hidden layer (see **Material and Methods**). Median was considered as the point posterior parameter estimation for all parameters. First column provides the average absolute error; second column shows the mean-squared error; third column shows the mean-squared error scaled by the parameter’s observed variance (see **Material and Methods** for error formulas).

| *AfrDE-EurDE*<br>*parameters* | ACB |  |  | ASW |  |  |
| --- | --- | --- | --- | --- | --- | --- |
|  | Av. absolute<br>Error | Mean-square<br>Error | Mean-square Error /<br>Var. | Av. absolute<br>Error | Mean-square<br>Error | Mean-square Error /<br>Var. |
| *S*Afr,0 | 0.2530 | 0.0857 | 1.0070 | 0.2444 | 0.0805 | 1.0081 |
| *S*Afr,1 | 0.1206 | 0.0216 | 0.8533 | 0.1158 | 0.0197 | 0.9259 |
| *S*Afr,20 | 0.02744 | 0.0012 | 0.4162 | 0.0219 | 0.0007 | 0.4773 |
| *u*Afr | 0.1166 | 0.0198 | 0.9974 | 0.1254 | 0.0216 | 0.9757 |
| *S*Eur,1 | 0.0952 | 0.0164 | 1.0526 | 0.1001 | 0.0157 | 1.0152 |
| *S*Eur,20 | 0.0044 | 0.0001 | 0.6452 | 0.0069 | 0.0001 | 0.6623 |
| *u*Eur | 0.1084 | 0.0174 | 0.9431 | 0.1021 | 0.0153 | 0.8036 |

Finally, based on these cross-validation procedures, we evaluated *a posteriori* if, in the vicinity of the ACB and ASW observed datasets respectively, the lengths of the estimated 95% credibility intervals for each parameter was accurately estimated or not (JAY *et al*. 2019). To do so, we calculated how many times the true parameter (*θ_i_*) was found inside the estimated 95% credibility interval [2.5%quantile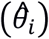; 97.5%quantile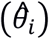], among the 1,000 out-of-bag NN-ABC posterior parameter estimation, separately for the ACB and ASW (**Supplementary Table S3**). For each parameter, if less than 95% of the true parameter values are found inside the 95% credibility interval estimated for the observed data, we consider the length of this credibility interval as underestimated indicative of a non-conservative behavior of the parameter estimation. Alternatively, if more than 95% of the true parameter-values are found inside the estimated 95% credibility interval, we consider its length as overestimated, indicative of an excessively conservative behavior of this parameter estimation.

#### Comparing the accuracy of posterior parameters estimations using NN, RF, or Rejection ABC

With the above procedure, we aimed at estimating the posterior parameter distributions jointly for all parameters, and their errors for the scenario most likely explaining observed genetic data for the ACB and ASW respectively. Nevertheless, NN-ABC and RF-ABC parameter inference procedures also allow estimating each parameter posterior distribution in turn and separately rather than jointly. This can further provide insights into how both ABC parameter inference approaches perform in the parameter space of the winning scenarios. To do so, we performed several out-of-bag cross-validation parameter estimation analyses for the ACB and ASW separately.

We compared four methods: NN estimation of the parameters taken jointly as a vector (similarly as in the above procedure), NN estimation of the parameters taken in turn separately, RF estimation of the parameters which also considers parameters in turn and separately (RAYNAL *et al*. 2019), and simple Rejection estimation for each parameter separately (PRITCHARD *et al*. 1999). For each method, we used in turn the 1,000 simulations closest to the real data as pseudo-observed data, and set a tolerance level of 1% of the 99,999 remaining simulations. We consider four neurons in the hidden-layer per neural network, and we considered 500 decision trees per random forest to limit the computational cost of these analyses at little accuracy cost *a priori* (**Supplementary Figure S6**). We then computed the mean-squared errors scaled by the variance of the true parameters, the mean-squared errors, and the mean absolute errors similarly as previously. Finally, we estimated the accuracy of the 95% credibility intervals for each method and for each parameter similarly as previously.

## RESULTS

First, we present results about the ability of *MetHis* to simulate data close to the observed ones. Second, we evaluate the ability of RF-ABC to distinguish, in the entire parameter space, the nine complex admixture scenarios in competition, and evaluate how each one of the 24 summary statistics contribute to distinguish among scenarios. Third, we use Random-Forest ABC to specifically predict the best fitting scenario for the history of admixture of two recently admixed populations descending from the Transatlantic Slave Trade in the Americas (African American ASW and Barbadian ACB). Fourth, we use Neural-Network ABC to estimate posterior parameter distributions under the winning scenario for the ACB and the ASW separately. Fifth, we evaluate in detail the accuracy of our posterior parameter estimation, and compare with other ABC posterior parameter inference approaches. Finally, we synthesize the complex admixture history thus reconstructed for the ASW and ACB populations.

### Simulating the observed data with MetHis

With *MetHis*, we conducted 10,000 simulations for each one of the nine competing scenarios for the admixture history of the ASW or the ACB populations, described in detail in **Figure 1** and **Material and Methods,** with corresponding model parameters drawn in *a priori* distributions described in **Table 1**.

We produced 90,000 vectors of 24 summary statistics each, overall highly consistent with the observed ones for the ACB and the ASW populations respectively. First, we found that each observed statistic is visually reasonably well simulated under the nine competing scenarios here considered (**Supplementary Figure S3**). Second, the observed data each fell into the simulated sets of summary statistics projected in the first four PCA dimensions (**Supplementary Figure S4**) considering all 24 summary statistics in the analysis. Finally, the observed vectors of 24 summary statistics computed for the ACB and ASW, respectively, were not significantly different (p-value = 0.468 and 0.710 respectively) from the 90,000 simulated sets of statistics using a goodness-of-fit approach (**Supplementary Figure S5**). Therefore, we successfully simulated datasets producing sets of summary statistics reasonably close to the observed ones, despite considering constant effective population sizes, fixed virtual source population genetic pool-sets, and neglecting mutation during the 21 generations of forward-in-time simulations performed using *MetHis*.

### Complex admixture scenarios cross-validation with RF-ABC

We trained the RF-ABC model-choice algorithm using 1,000 trees, which guaranteed the convergence of the model-choice prior error rates (**Supplementary Figure S6**). Based on this training, the complete out-of-bag cross-validation matrix showed that the nine competing scenarios of complex historical admixture could be relatively reasonably distinguished using our set of 24 summary statistics and 10,000 simulations under each competing scenario, despite the high level of nestedness of the scenarios here considered (see **Material and Methods**). Indeed, we calculated an out-of-bag prior error rate of 32.41%, considering each 90,000 simulation, in turn, as out-of-bag pseudo-observed target dataset and the rest of simulations (89,999) as the training dataset for RF-ABC scenario-choice. Furthermore, we found the posterior probabilities of identifying the correct scenario ranging from 55.17% (prior probability = 11.11% for each competing scenario), for the two-pulses scenarios from both the African and European sources (Afr2P-Eur2P), to 77.71% for the scenarios considering monotonically decreasing recurring admixture from both sources (AfrDE-EurDE) (**Figure 2A**).

Importantly, the average probability, for a given admixture scenario, of choosing any one alternative (wrong) scenario were on average 4.05% across the eight alternative scenarios, ranging from 2.79% for the AfrDE-EurDE scenario, to 5.60% for the Afr2P-Eur2P scenario (**Figure 2A**). This shows that our approach did not systematically favor one or the other competing scenario when wrongly choosing a scenario instead of the true one, despite high levels of nestedness among scenarios.

We find that the six summary statistics most contributing to the observed cross validation results for RF-ABC model-choice among the 24 statistics here tested were statistics describing specifically the admixture-fraction distribution: minimum and maximum admixture fraction values, variance, skewness, as well as the 10% and 90% quantiles of the distribution (**Figure 2B**). Interestingly, within and between populations summary-statistics often used in population genetics (including *F_ST_*, mean heterozygosity, and *f3* statistics), contributed to distinguishing the competing complex admixture scenarios to a lesser extent.

Finally, note that scenarios considering monotonically recurring admixture from each source populations (AfrDE-EurDE, AfrDE-EurIN, AfrIN-EurDE, AfrIN-EurIN) can be relatively well distinguished, using our RF-ABC framework, from scenarios with at least one source population contributing to the admixed population with two possible pulses after the foundation event (Afr2P-Eur2P, Afr2P-EurDE, Afr2P-EurIN, AfrDE-Eur2P, AfrIN-Eur2P). Indeed, we found an out-of-bag prior error rate of 13.85%, and posterior cross-validation probabilities of identifying the correct group of scenarios of 86.08% and 86.23% respectively for the two groups (ESTOUP *et al*. 2018).

### Complex admixture histories for the Barbadian and African American populations

#### Random-Forest ABC scenario-choice

We performed RF-ABC model-choice with 1,000 decision trees and 10,000 simulations per each nine competing scenarios (**Figure 1** and **Table 1**, **Material and Methods**), separately for the admixture history of the Barbadian (ACB) and the African American (ASW) populations. For the ACB, **Figure 3** shows that the majority of votes (53.1%) went to an admixture scenario AfrDE-EurDE with a posterior probability of the winning scenario of 60.28%. This scenario encompassed monotonically decreasing recurring contributions from both the African and European source populations over the last 20 generations before present. The second most chosen scenario considered a monotonically decreasing recurring contribution from the African source population over the last 20 generations, while the European source population contributed two admixture pulses to this admixed population after the founding pulse (scenario AfrDE-Eur2P). However, this scenario is voted for 3.5 times less often than the winning scenario AfrDE-EurDE, gathering 15.1% of the 1,000 votes, only slightly above the 11.11% prior probability for each nine-competing scenario (**Figure 3**; **Supplementary Table S1**).

Concerning the admixture history of the ASW, RF-ABC scenario-choice results were less segregating. **Figure 3** shows that the AfrDE-EurDE scenario also gathered the majority of votes for the admixture history of the ASW, albeit with lower posterior probability than for the ACB (33.5% of 1,000 votes, with posterior probability = 48.0% for the ASW). The second most chosen scenario, AfrDE-Eur2P, was only slightly less chosen with 31.7% of the votes (**Figure 3**, **Supplementary Table S1**). For the ASW, considering only the two best scenarios (AfrDE-EurDE and AfrDE-Eur2P) to train the Random Forest, and re-conducting the RF-ABC scenario-choice, improved the scenario discrimination in favor of the AfrDE-EurDE scenario. While we found only a slight majority of votes (51.8%) also in favor of the AfrDE-EurDE scenario, we found a substantially increased posterior probability for this model equal to 57.9%. This increased posterior probability of the AfrDE-EurDE scenario compared to the previous RF-ABC scenario-choice considering the nine competing scenarios (48.0%), indicated that this scenario best explains the ASW observed genetic patterns, despite overall limited discriminatory power of our approach in the part of the summary-statistics space occupied by the ASW.

#### Neural-Network ABC parameter inference accuracy for the ACB and ASW populations

We performed 100,000 simulations using *MetHis* for the AfrDE-EurDE scenarios, in order to estimate, using Neural-Network ABC, posterior parameter distributions and the corresponding parameter prediction cross-validation errors, considering in turn the ACB and the ASW populations (**Figure 4 and 5**, **Table 2**, **Table 3**, and **Supplementary Table S3**).

**Figure 5:**
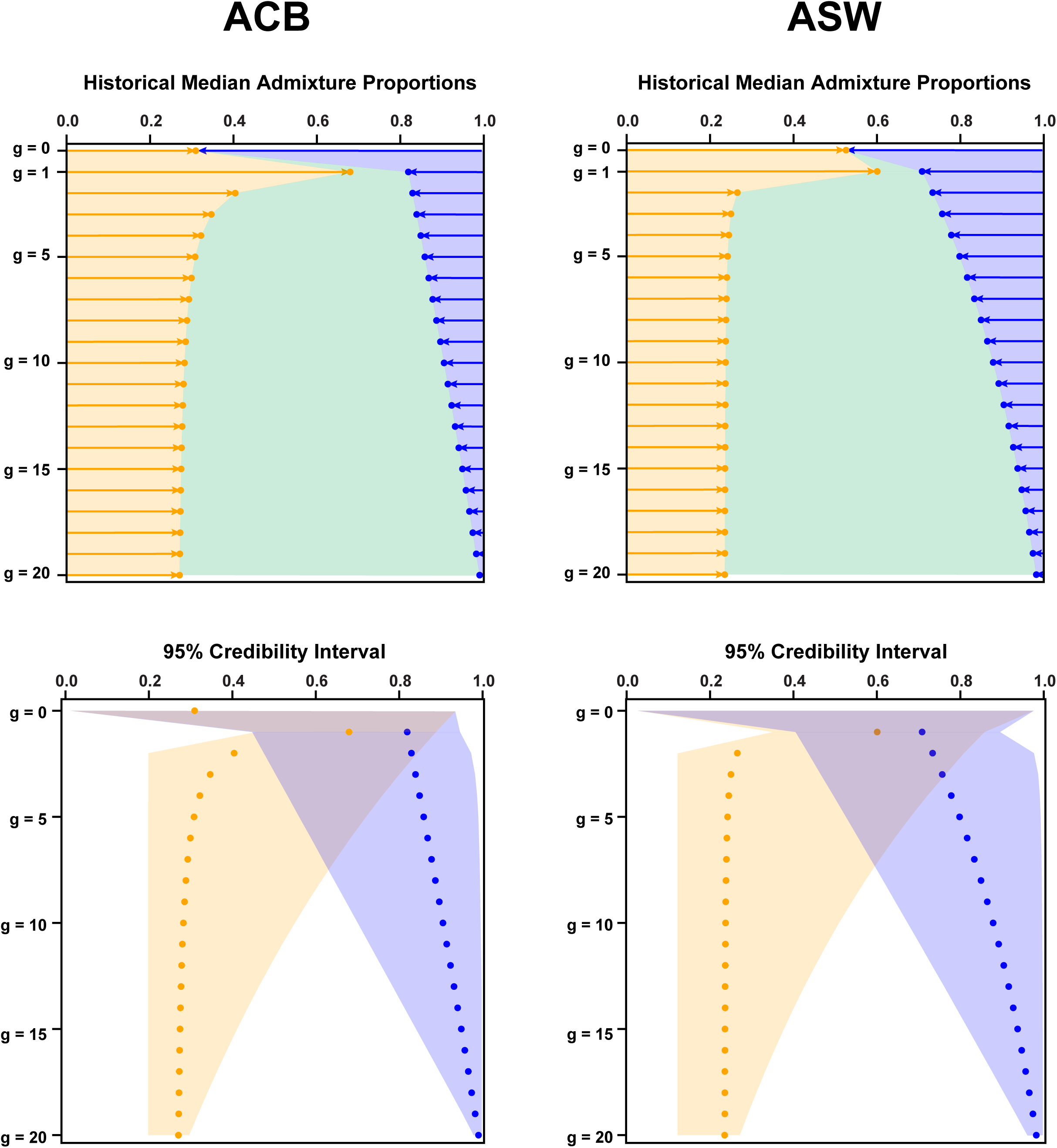
Approximate Bayesian Computation inference of the admixture history of the ACB and ASW populations respectively. Top panels are based on median point-estimates of intensity parameters at each generation. Bottom panels show 95% credibility intervals for each inferred parameter around the median point-estimates. The African introgression is plotted in orange, the European introgression in blue, and in green the remaining contribution of the admixed population to itself at the following generation. (A) Results for the ACB under the AfrDE-EurDE winning scenario; (B) Results for the ASW under the AfrDE-EurDE winning scenario.

For the ACB under the AfrDE-EurDE scenario (**Figure 4A**, **Table 2**), we found that the two recent admixture intensities from Africa and Europe (*s*_Afr,20_ and *s*_Eur,20_, respectively) and the steepness of the European decrease in contribution over time (*u*_Eur_) had sharp posterior densities clearly distinct from their respective priors. Note that the cross-validation error on these parameters in the vicinity of our real data were low (average absolute error 0.02744, 0.0044, and 0.1084, respectively for *s*_Afr,20_, *s*_Eur,20_, and *u*_Eur_) (**Table 3**), and lengths of 95% credibility intervals reasonably accurate (96.4%, 94.4%, 94.1% of 1,000 cross-validation true parameter values fell into estimated 95% credibility intervals, **Supplementary Table S3**). This shows the reliability of our method to accurately infer the three parameters in the part of the space of summary statistics occupied by the ACB observed data.

Furthermore, the two ancient admixture intensities from Africa and Europe at generation 1 immediately following the initial foundation of the admixed population H (*s*_Afr,1_ and *s*_Eur,1_, respectively), also had posterior densities apparently distinguished from their prior distributions, but both had much wider 95% credibility intervals (**Figure 4A**, **Table 2**). Consistently, we found a slightly increased posterior parameter error in this part of the parameter space for both these parameters, with average absolute error 0.121 and 0.095 respectively for *s*_Afr,1_ and *s*_Eur,1_ (**Table 3**). Nevertheless, note that 95.8% and 94.7% of 1,000 cross-validation true values for those two parameters fell into the estimated 95% credibility intervals (**Supplementary Table S3)**. This shows a reasonably conservative behavior of our method for these estimations, further indicating that information is lacking in our data or set of summary statistics for a more accurate estimation of these parameters, rather than an inaccuracy of our approach.

Interestingly (**Figure 4A**, **Table 2**), we found that accurate posterior estimation of the steepness of the African decrease in admixture over time (*u*_Afr_) is difficult. Indeed, the posterior density of this parameter only showed a tendency towards small values slightly departing from the prior, indicative of a limit of our method to estimate this parameter (**Figure 4A**, **Table 2**). Finally (**Figure 4A**, **Table 2**), we found that we had virtually no information to estimate the founding admixture proportions from Africa and Europe at generation 0, as our posterior estimates barely departed from the prior and associated mean absolute error was high (0.2530, **Table 3**). Nevertheless, our method seemed to be performing reasonably conservatively for these two latter parameters (95.6% and 95.3% of 1,000 cross-validation true parameter values fell into estimated 95% credibility intervals, **Supplementary Table S3**). This indicates that information is strongly lacking in our data or summary statistics for successfully capturing these parameters, rather than inherent inaccuracy of our ABC method.

For the African American ASW under the AfrDE-EurDE model, our posterior parameter estimation accuracy results were overall quantitatively slightly less accurately estimated compared to those obtained for the ACB population, as indicated by overall larger credibility intervals and cross-validation errors (**Figure 4B**, **Table 2**, **Table 3**, **Supplementary Table S3**). This was consistent with the more ambiguous RF-ABC model-choice results obtained for this population (**Figure 3**).

#### Comparing NN, RF, and Rejection ABC posterior parameter estimation accuracy

For posterior parameter estimations considering the ACB or the ASW population, the means of the three types of errors (scaled mean-square error, mean-square error, absolute error, see **Material and Methods**) were systematically lower for the two NN methods (joint or independent posterior parameter estimation) than for the RF and Rejection independent posterior parameter estimation methods (**Table 4**). Furthermore, we found that the means of the three types of errors were qualitatively comparable between the NN estimation of the parameters taken as a joint vector and the NN estimation of the parameters taken separately. Altogether, these results showed that considering the NN estimation for parameters taken jointly as a vector is overall preferable for the ACB and ASW populations, since it further allowed the joint interpretation of parameter values estimated *a posteriori*, with little difference in accuracy between the two methods.

**Table 4.** Approximate Bayesian Computation mean posterior parameter errors under the winning Scenario AfrDE-EurDE, for the ACB and ASW populations separately, using four different methods: NN estimation of the parameters taken jointly as a vector, NN estimation of the parameters taken separately, Random Forest (parameters taken separately), and Rejection (parameters taken separately). For each target population separately and for each method, we conducted an out-of-bag cross validation by considering in turn 1,000 separate parameter inferences each using one of the 1,000 closest simulation to the observed ACB (or ASW) data as the target pseudo-observed dataset. All posterior parameter estimations were conducted using the other 99,999 simulations under the AfrDE-EurDE scenario (**Figure 1**, Table 1), a 1% tolerance rate (i.e. 1,000 simulations), 24 summary statistics, logit transformation of all parameters, four neurons in the hidden layer per neural network and 500 trees per random forest. Median was considered as the point posterior parameter estimation for all parameters. First column provides the average absolute error; second column shows the mean-squared error; third column shows the mean-squared error scaled by the parameter’s observed variance (see **Material and Methods** for error formulas).

| *Posterior parameter*<br>*estimation ABC*<br>*method* | ACB |  |  | ASW |  |  |
| --- | --- | --- | --- | --- | --- | --- |
|  | Av. absolute<br>Error | Mean-squared<br>Error | Mean-squared Error<br>/ Var. | Av. absolute<br>Error | Mean-squared<br>Error | Mean-squared Error<br>/ Var. |
| *NN joint* | 0.1037 | 0.0232 | 0.8450 | 0.1024 | 0.0219 | 0.8383 |
| *NN independent* | 0.1032 | 0.0236 | 0.8294 | 0.1025 | 0.0225 | 0.8344 |
| *RF independent* | 0.1042 | 0.0246 | 0.8534 | 0.1036 | 0.0233 | 0.8697 |
| *Rejection*<br>*independent* | 0.1071 | 0.0238 | 0.9299 | 0.1050 | 0.0223 | 0.8951 |

Finally, results showed that the lengths of 95% credibility intervals estimated with NN joint parameter estimation was, across all parameters, more accurate than all other methods with, on average, 95.1% and 95.2% of true parameter values falling within the estimated 95% credibility intervals, for the ACB and ASW respectively (**Supplementary Table S3**). Furthermore, we found that lengths of 95% credibility intervals estimated with NN and RF independent posterior parameter estimations were systematically under-estimated, with less than 94% of the true parameter values falling into the 95% credibility intervals estimated. Finally, we found that lengths of 95% credibility intervals estimated with the Rejection method were also rather accurately estimated although on average slightly over-estimated compared to the NN joint parameter estimation with, on average, 95.5% of the 1,000 cross-validation true parameter values within the estimated 95% credibility intervals for the ACB, and 95.8% for the ASW.

#### Admixture histories of the African American ASW and Barbadian ACB

**Figure 5** visually synthesized the estimated posterior parameters of the complex admixture scenarios reconstructed with our novel *MetHis* – machine-learning ABC framework, and associated 95% credibility intervals (**Table 2**).

We found a virtual complete replacement of the ACB and ASW populations at generation 1 after foundation, thus consistent with our inability to accurately estimate the founding proportions from the African and European sources at generation 0. Furthermore, we found an increasingly precise posterior estimation of African and European contributions to the gene-pool of the ACB and ASW populations forward in time, with most recent estimations exhibiting narrow credibility intervals. This is also consistent with the nature of recurrent admixture processes, where older information is often lost or replaced when more recent admixture events occur.

Most interestingly, we found that the recurring contribution of the European gene pool to the admixed populations rapidly decreases after generation 1 for both the ACB and ASW albeit with substantial differences (**Figure 5**). Indeed, we found that the recurring contribution from the European source to the ACB gene pool falls below 10% at generation 9 until no more than 1% in the present (generation 20). Comparatively, we found that the European contribution diminished substantially less rapidly for the ASW, going below 10% only after generation 12 until roughly 2% in the present. This indicates that the European contribution to the African American gene pool was more sustained over time than for the Barbadian.

Finally, we found substantial recurring contributions from the African source population to the gene pool of both admixed populations (**Figure 5**). For the ACB population, we found a progressive decrease of the African recurring introgression until a virtually constant recurring admixture close to 28% from generation 10 and onward. For the ASW, our results showed a sharper decrease of the African contribution after foundation until a virtually constant recurring admixture process close to 24% from generation 5 until present (generation 20). The high overall African recurring introgression into the admixed-populations gene pools captures the importance of recurring admixture in explaining the observed patterns for both populations descending from the TAST.

## DISCUSSION

We evaluated how machine-learning Approximate Bayesian Computation methods can bring new insights to the reconstruction of highly complex admixture histories using genetic data. To illustrate our proof of concept and thoroughly investigate the power and accuracy of our approach using real data, we aimed at reconstructing the recent complex admixture history for the African American (ASW) and Barbadian (ACB) population samples from the 1000 Genomes project (Phase 3).

Our results demonstrated that our novel *MetHis* forward-in-time simulator and summary statistics calculator coupled with RF-ABC scenario-choice can often clearly infer the best class of highly complex admixture histories underlying independent SNP data diversity, in a reasonable-size sample and genetic dataset. In the two source-populations admixture models here investigated, we distinguished scenarios encompassing two pulses of admixture from each source, after the founding admixture event, monotonically increasing or decreasing admixture intensities over time, or a combination of these three scenarios. Furthermore, we found that NN-ABC provide accurate posterior parameter inference of most demographic parameters of recurring monotonically decreasing admixture processes, compared to other classes of ABC posterior parameter inference methods. Finally, we empirically demonstrated that the moments of the distribution of admixture fractions within the admixed population estimated using independent SNPs were highly informative for reconstructing the admixture history using an ABC approach, as expected theoretically (VERDU and ROSENBERG 2011; GRAVEL 2012).

While we found that distinguishing among competing models is more difficult in certain parts of the parameter space due to scenario-nestedness (ROBERT *et al*. 2010), our *MetHis* – ABC method already vastly extends the array of complex admixture models explored with most, classically used, maximum-likelihood inference approaches (ROBERT *et al*. 2010; GRAVEL 2012; LOH *et al*. 2013; HELLENTHAL *et al*. 2014). It is challenging to analytically predict genomic diversity patterns expected under realistic complex admixture histories, as likelihood calculations under such models are very often intractable (VERDU and ROSENBERG 2011; GRAVEL 2012; MEDINA *et al*. 2018; NI *et al*. 2019). In turn, this makes it difficult to understand how most existing efficient maximum-likelihood admixture inference methods, which often only consider one or two pulses of admixture, behave when the observed genetic data in fact results from much more complex admixture processes (GRAVEL 2012; HELLENTHAL *et al*. 2014).

In this context, the proof of concept here presented more generally shows that ABC can be fruitfully attempted to explore, virtually, any other admixture model beyond the case studies here-conducted, provided that, *a priori*, simulation and summary statistics calculation are feasible. To these ends, other recent efficient forward-in-time genetic data simulators can also be successfully used in an ABC framework instead of *MetHis* (HALLER and MESSER 2019; NI *et al*. 2019). In reality, studies investigating ABC approaches for admixture reconstruction, while allowing for exploring scenarios out of reach of other methods, will inevitably face the same difficulties as any ABC inference; such as high dimensional parameter and summary-statistics spaces, lack of information from summary statistics, and scenario nestedness (CSILLÉRY *et al*. 2010; ROBERT *et al*. 2010; SISSON *et al*. 2018).

Importantly, the current *MetHis* – ABC approach does not make use of admixture linkage-disequilibrium patterns in the admixed population, and only relies on independent genetic markers. Nevertheless, admixture LD has consistently proved to bring massive information about the complex admixture history of numerous populations worldwide (GRAVEL 2012; HELLENTHAL *et al*. 2014; MEDINA *et al*. 2018; NI *et al*. 2019). However, existing methods to calculate admixture LD patterns remain computationally intensive and require numerous markers and accurate phasing, which is difficult under ABC where such statistics have to be calculated for each one of the numerous simulated datasets. In this context, RF-ABC (PUDLO *et al*. 2016; RAYNAL *et al*. 2019) or AABC (BUZBAS and ROSENBERG 2015) methods allow substantially diminishing the number of simulations required for satisfactory scenario-choice and posterior parameter inference, which makes both approaches promising tools for using, in the future, admixture LD patterns to reconstruct complex admixture processes from genomic data.

Sex-biased admixture processes are known to have influenced admixed populations, and in particular populations descending from the TAST (MORENO-ESTRADA *et al*. 2013; FORTES-LIMA *et al*. 2018). Future version of our *MetHis* – ABC framework will explicitly implement sex-specific admixture processes with, in addition to autosomal data, the possibility to investigate sex-related genetic data (X-chromosome, Y-chromosome, and mitochondrial DNA) (GOLDBERG *et al*. 2014; GOLDBERG and ROSENBERG 2015).

Finally, although *MetHis* readily allows considering changes of effective population size in the admixed population at each generation as a parameter of interest to ABC inference, we did not, for simplicity, investigate here how such changes affected our results for the African American and Barbadian admixed population. Future work using *MetHis* will allow specifically investigating how effective size changes may influence genetic patterns in the admixed population, a question of major interest as numerous admixed populations are expected to have experienced founding events and/or bottlenecks during their history (e.g. BROWNING *et al*. 2018).

For all these reasons, it is crucial, in general and in the future, to further develop novel methodological tools and evaluate how genetic patterns evolve over time as a function of each parameter of complex historical admixture models separately (BUZBAS and VERDU 2018; MEDINA *et al*. 2018; NI *et al*. 2019). *MetHis* can help to this task since it allows the users to investigate how parameters of the complex admixture process can influence, over time, a large number of population genetics summary-statistics calculated in the simulated admixed population at each generation.

Concerning the specific admixture history of the two admixed populations descending from the TAST here reconstructed, note that several competing scenarios can clearly be discarded for explaining the observed genetic patterns. In particular, the Afr2P – Eur2P scenario considering two possible pulses of introgression after the founding event, separately from the African and European source, does not significantly exceeds the prior probability of choosing any nine-competing scenario (4.6% and 11.2% of the 1,000 votes for the Afr2P – Eur2P, respectively for the ACB and the ASW, **Figure 3**). Note that this scenario embeds models analogous to the most complex admixture scenarios that have been previously tested for these populations with maximum-likelihood approaches based on extensive genome-wide data and admixture-LD based statistics (GRAVEL 2012; BAHARIAN *et al*. 2016). Interestingly, very recent migrations from either Africa or Europe to the Americas are known to have been intense demographically in the 19^th^ and 20^th^ century (BERLIN 2010). However, the recent increased demographic migrations do not seem to have left the equivalent signature in the genetic admixture process of both the ACB and ASW populations, as monotonically recurring increasing admixture scenarios can here be rejected confidently.

Nevertheless, we found that genetic admixture of African origin in both admixed populations, although decreasing since foundation, retained high levels in the present day (between 20% and 30%). These results could stem from the known importance of African recurring forced migrations during the TAST into the Americas; further prompts the influence of African slave descendants forced migrations within the Americas after the initial crossing of the ocean (often called the Middle Passage); and highlights the major importance of post-slavery migrations of TAST descendant populations within the Americas (BERLIN 2010; ELTIS and RICHARDSON 2010; BAHARIAN *et al*. 2016). For instance, intense migrations from Haitian slave-descendants in the 19^th^ century have already been shown to possibly have contributed to the admixture patterns of other populations in the Caribbean and continental America (MORENO-ESTRADA *et al*. 2013; FORTES-LIMA *et al*. 2018).

Finally, we found that the genetic contribution from Europe rapidly decreases, after the foundation of both admixed populations, to marginal amounts during the 20^th^ century. Therefore, it seems that neither sustained European migrations, nor the relaxation of social and legal constraints on admixture between descendant communities subsequent to the abolition of slavery and the end of segregation, have translated into increased European genetic contribution to the gene-pool of admixed populations descending from European and African forced or voluntary migrations into the Americas after the TAST.

Altogether, our results for the two recently-admixed human populations illustrated how our *MetHis* – ABC framework can bring fundamental new insights into the complex demographic history of admixed populations; a framework that can easily be adapted for investigating admixture history in numerous populations and species, particularly when maximum-likelihood methods are intractable.

## Acknowledgements

We warmly thank Frédéric Austerlitz, Erkan O. Buzbas, Antoine Cools, Flora Jay, Evelyne Heyer, Margueritte Lapierre, Guillaume Laval, Nina Marchi, Etienne Patin, Noah A. Rosenberg, and Zachary A. Szpiech for useful comments and discussions. This project was funded in part by the French Agence Nationale de la Recherche project METHIS (ANR 15-CE32-0009-01). CFL was funded in part by the Sven and Lilly Lawski’s Foundation (N2019-0040).

## Supplementary note S1

We used the rectangular hyperbola class of functions to obtain increasing/decreasing patterns using only one shape parameter. We give here the derivation of the equations used, giving the example of a decreasing pattern.

A decreasing hyperbola is given by the function:

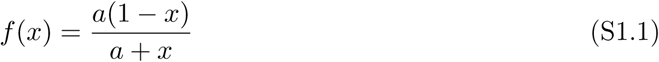

with *x ∈* [0; 1], *f* (*x*) *∈* [0; 1] and *a ∈* [0; +*∞*[. Parameter *a* controls the shape (“steepness”) of the curve obtained (figure S1.1).

**Figure S1.1:**
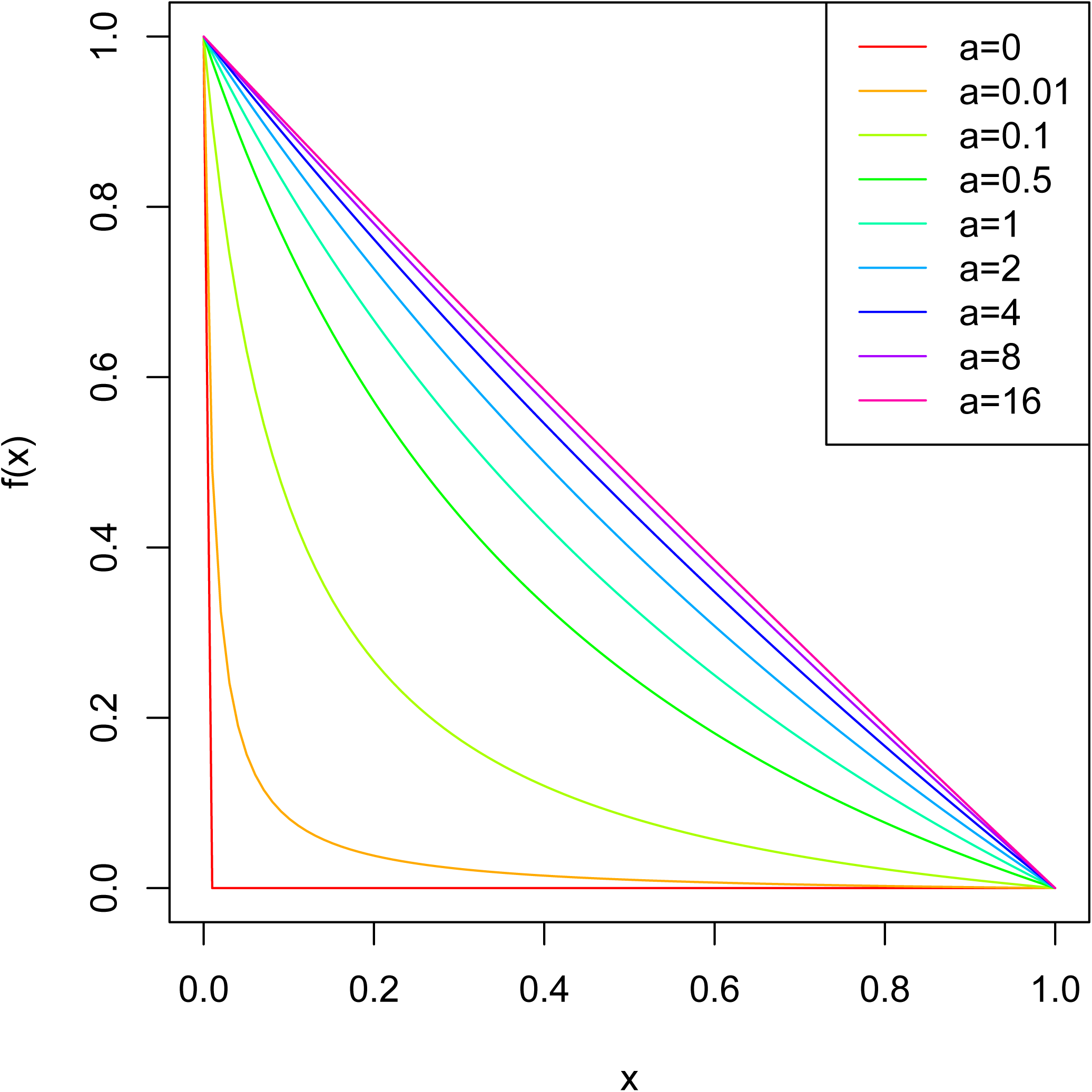
Influence of *a* on equation S1.1

The intersection between the hyperbola and *y* = *x* is given by

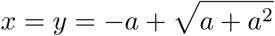

thus, we can sample an uniform deviate *u ∈* [0; 1/2] and set parameter *a*:

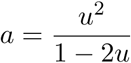

to obtain all hyperbola shapes.

We then transformed equation S1.1 to rescale the ranges of *x* and *f* (*x*):

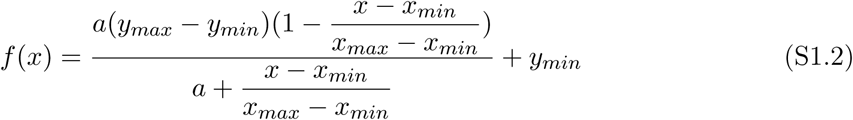

with *x ∈* [*x_min_*; *x_max_*] and *f* (*x*) *∈* [*y_min_*; *y_max_*] (figure S1.2).

**Figure S1.2:**
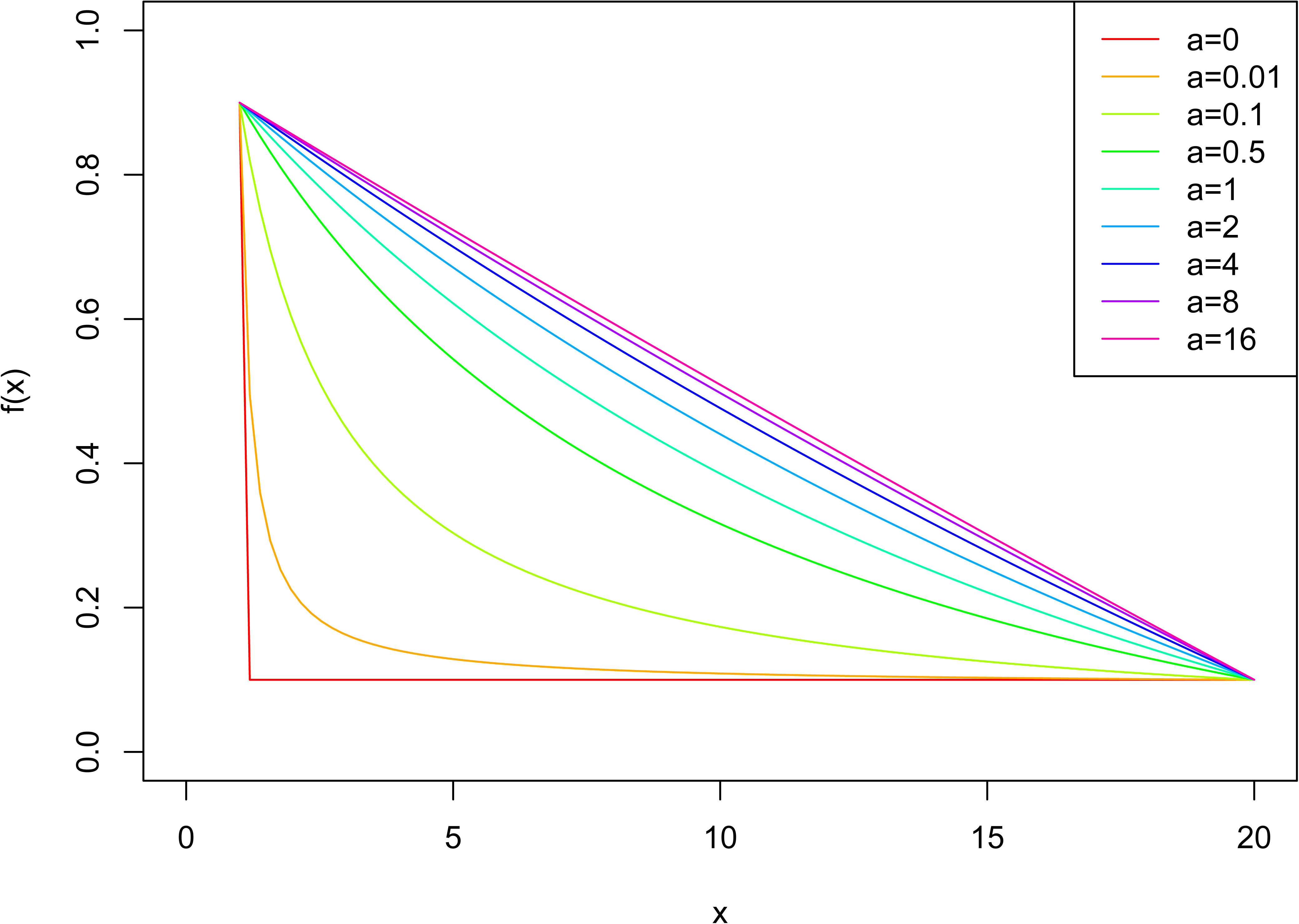
Influence of *a* on equation S1.2 with *x_min_* = 1, *x_max_* = 20, *y_min_* = 0.1 and *y_max_* = 0.9

With the notation used in the main text for contributions, and considering 20 generations of admixture, we obtain:

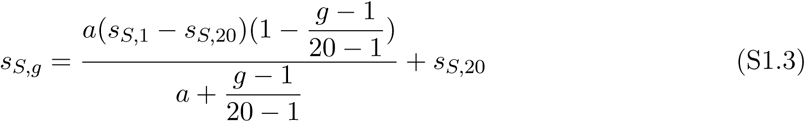

and an example of the patterns obtained for different *u* values is given in figure S1.3.

**Figure S1.3:**
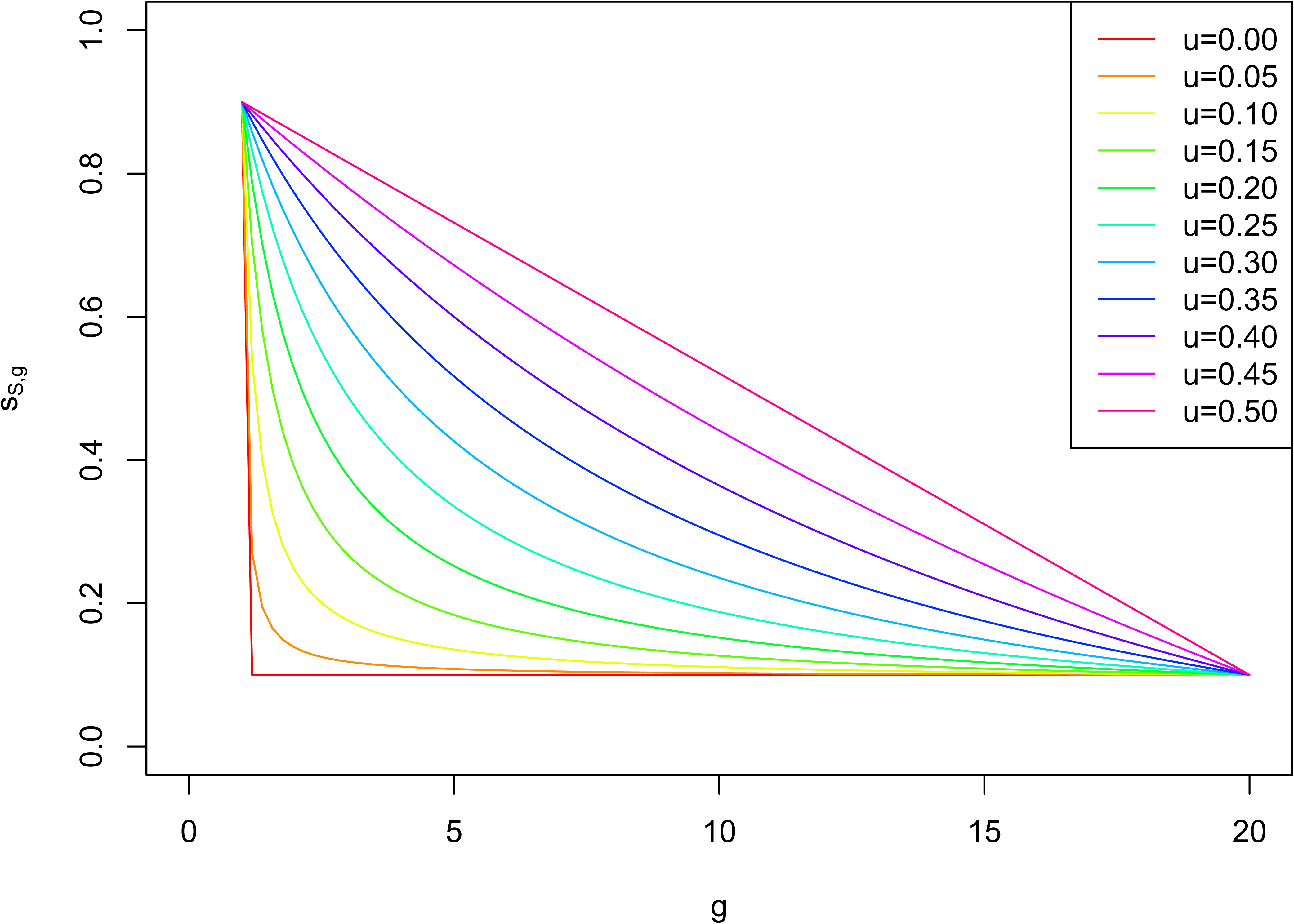
Influence of *u* on equation S1.3 with *s_S,_*_20_ = 0.1 and *s_S,_*_1_ = 0.9

---

**Supplementary Figure S1.**
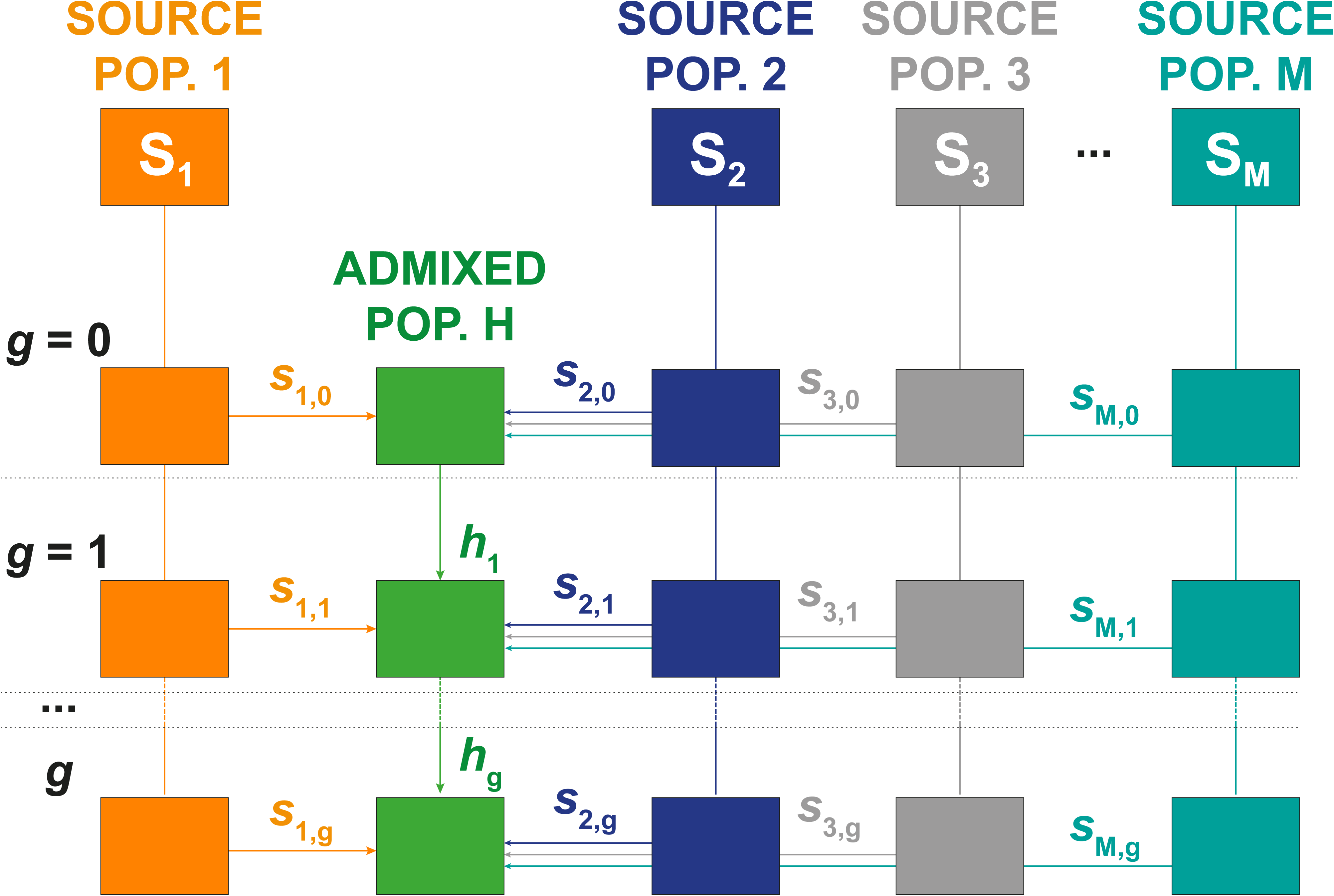
General mechanistic model of historical admixture from Verdu and Rosenberg (2011).

**Supplementary Figure S2:**
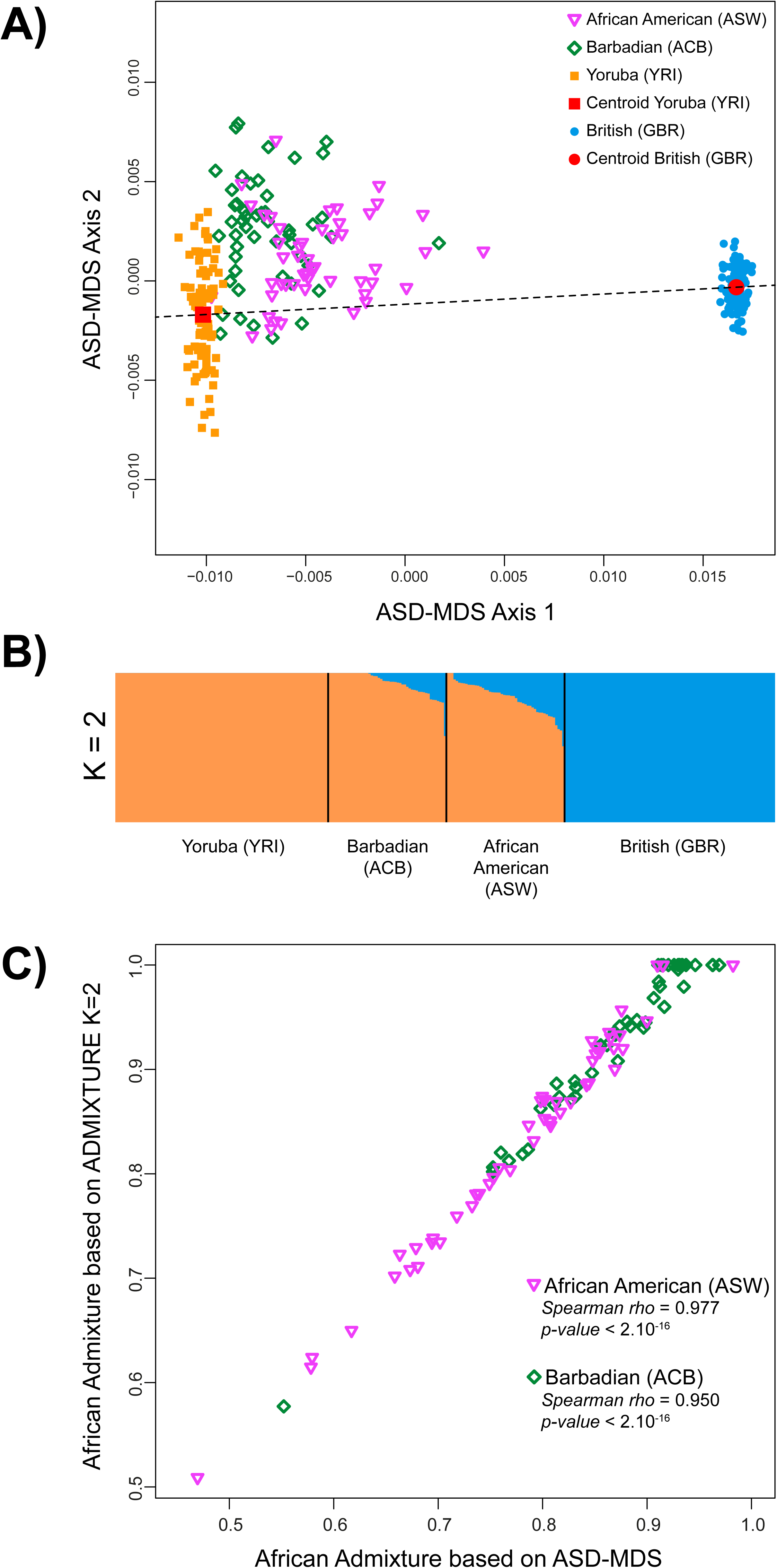
Comparison of individual admixture estimates using ASD-MDS and ADMIXTURE for the Barbadian (ACB) and the African American (ASW). 100,000 independent SNPs were considered from the 1000 Genome Project Phase 3 for 279 unrelated individuals (90 Yoruba (YRI), 89 British (GBR), 50 Barbadian (ACB), 50 African American (ASW)). (A) Allele Sharing Dissimilarity was computed between all pairs of individuals and the resulting matrix projected on the first two dimensions of a metric MDS. The two-dimensional centroid of the Yoruba (YRI) and, respectively, the British (GBR) are indicated in red and connected by a black dotted line. ACB and ASW individuals are projected orthogonally onto this line and their relative distance to the Yoruba centroid is calculated to obtain ASD-MDS based individual admixture estimates. (B) A single run of unsupervised ADMIXTURE (Alexander et al. 2009) has been computed using the 279 individuals and 100,000 SNPs and results were plotted using DISTRUCT (Rosenberg 2004). Individual membership proportions to the “orange” cluster mostly represented by Yoruba (YRI) genotypes was considered as an estimate of African admixture for the ACB and ASW respectively. (C) Spearman correlation between ASD-MDS and ADMIXTURE-based estimates of African admixture for the ACB and ASW individuals separately.

**Supplementary Figure S3:**
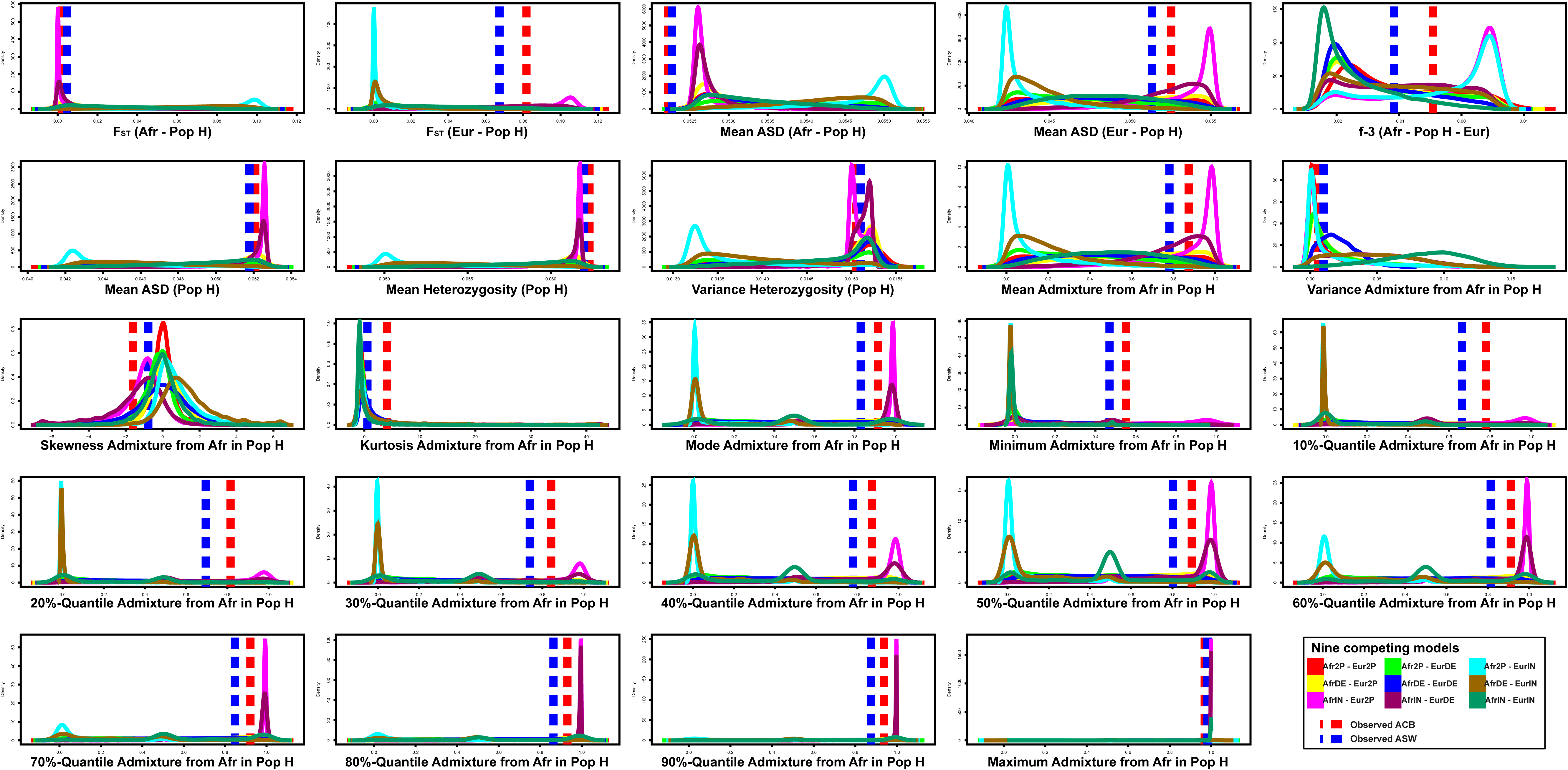
Summary statistics prior-distribution densities for each nine competing models considered (Figure 1). 10,000 simulations were performed for each nine-competing scenario and prior densities plotted with a different color indicated for each scenario. Corresponding statistics observed from the ACB and ASW population separately are represented, on each plot, by vertical doted-lines (red and blue respectively for ACB and ASW). The 24 separate summary statistics considered are described in **Material and Methods**.

**Supplementary Figure S4:**
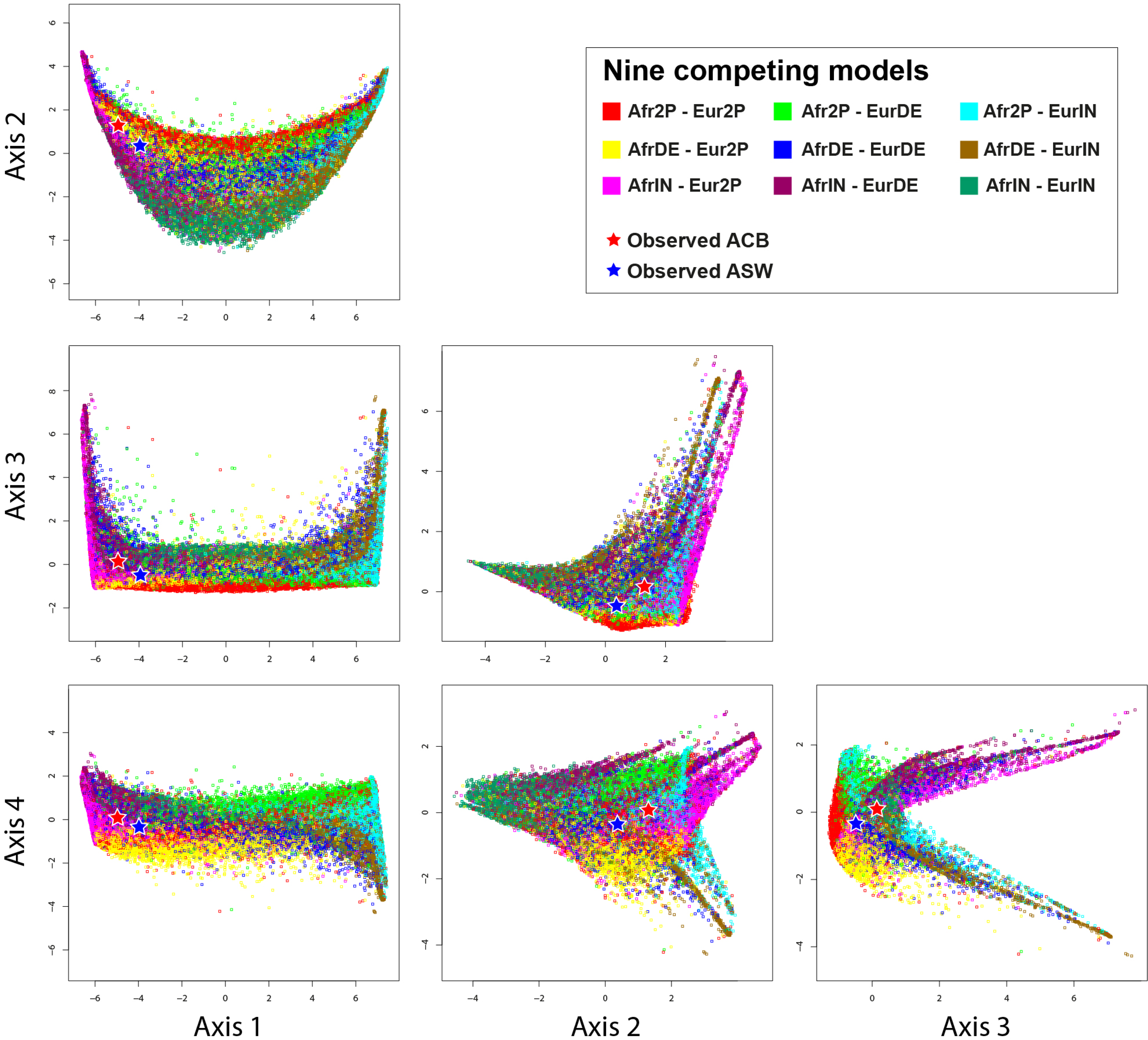
Four first axes of the principal component analysis for the 90,000 sets of 24 summary statistics computed on simulated data under each nine-competing scenario (Figure 1). The 24 same statistics calculated for the observed ACB and ASW population samples, respectively, are then projected on the PCA and represented by, respectively, a red and blue star. All two-dimensional projections are orthonormal.

**Supplementary Figure S5:**
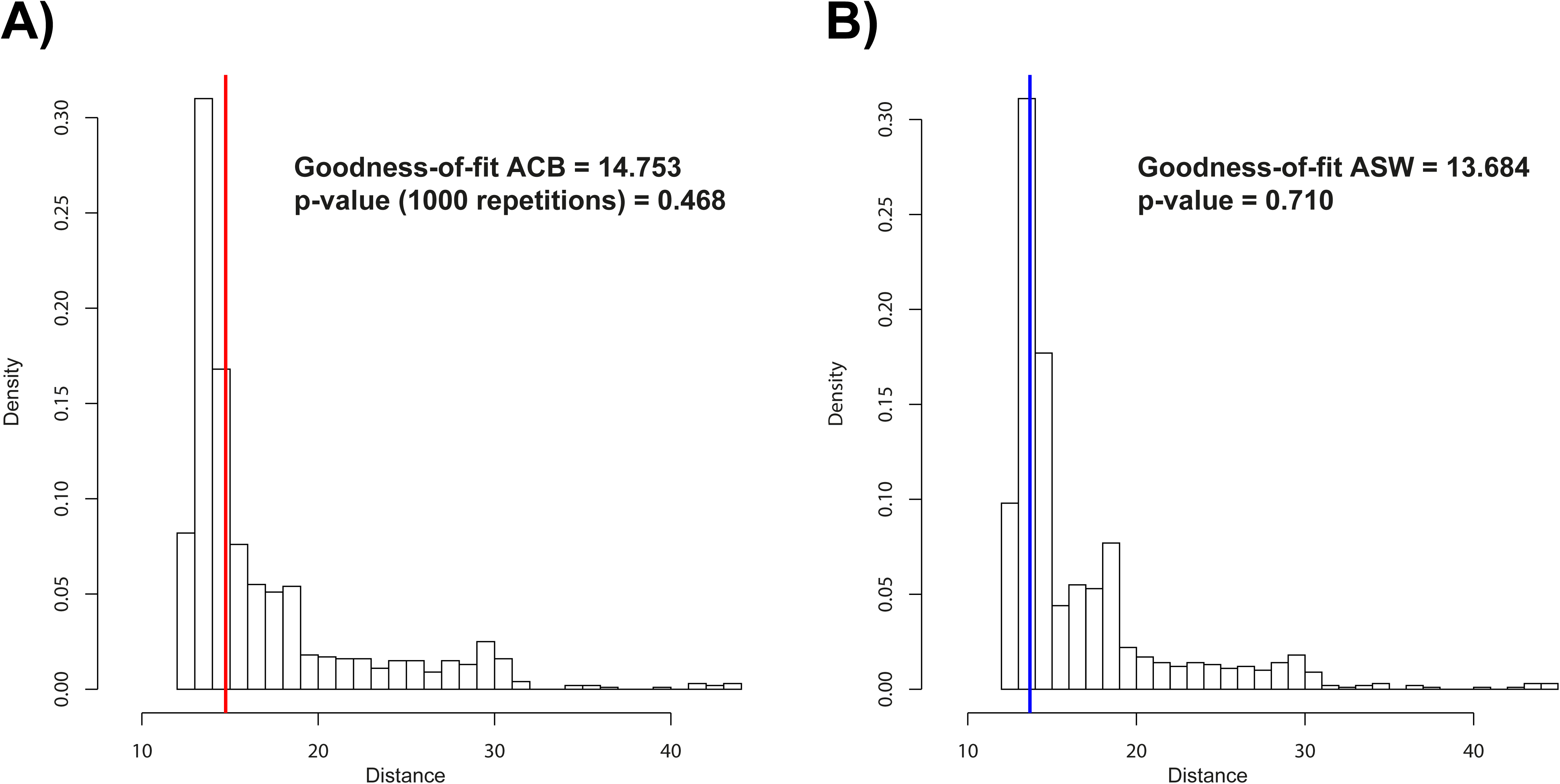
Histogram of the goodness-of-fit for the observed set of 24 summary statistics computed for (A) the ACB population, and (B) the ASW population, in turn serving as the observed admixed population H considering the YRI population sample as the African source and the GBR population sample as the European source (see **Material and Methods**). Goodness-of-fit statistics were calculated as the mean distance between observed and accepted summary statistics. Observed statistics are fitted to the full 90,000 sets of the same statistics calculated from 10,000 simulations performed under each nine-competing models (Figure 1). Goodness-of-fit was obtained considering 1,000 repetitions and a tolerance value of 0.01.

**Supplementary Figure S6:**
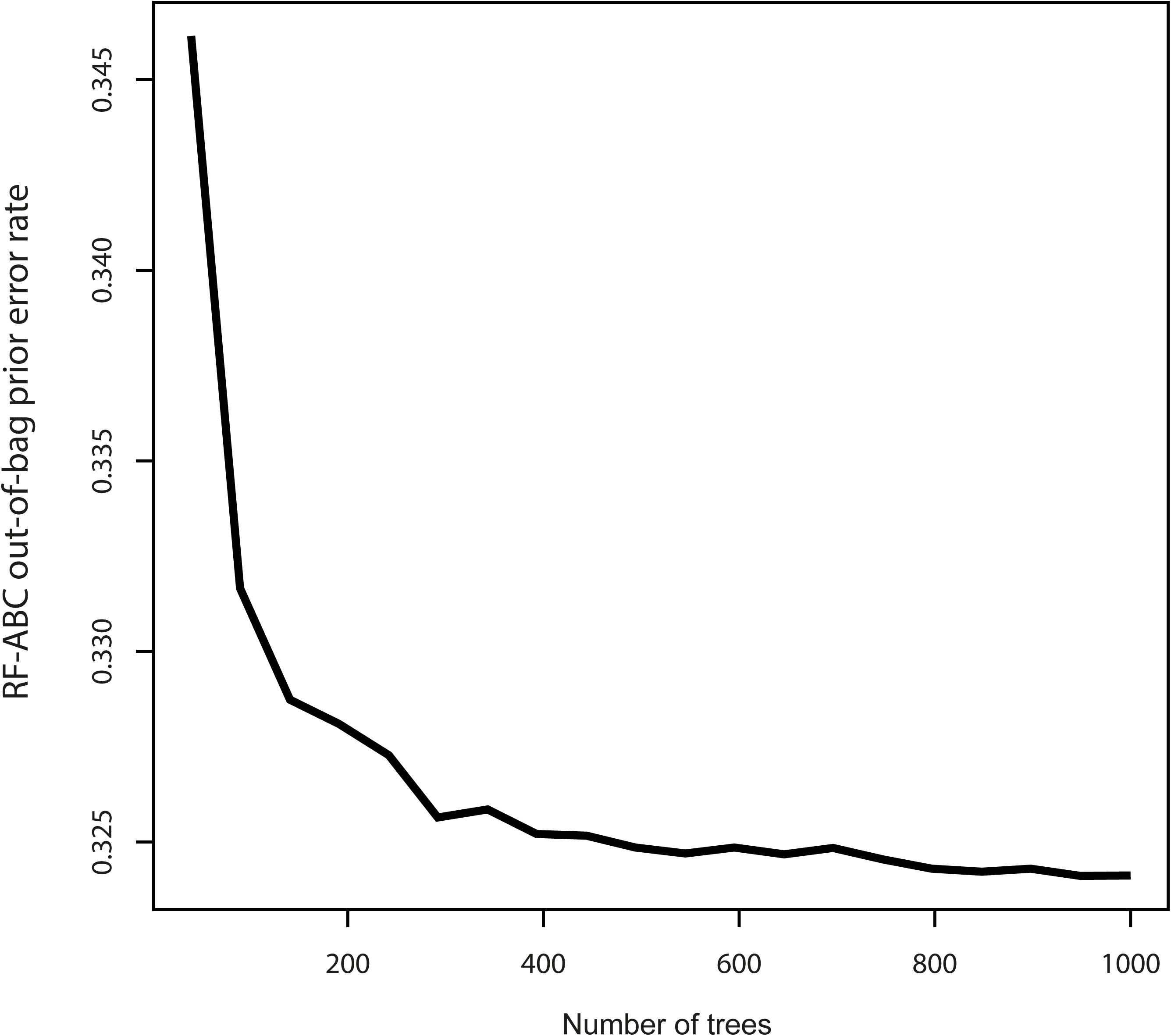
RF-ABC out-of-bag prior error rate as a function of the number of trees considered to build the forest for the model-choice procedure considering nine-competing scenarios (Figure 1).

**Supplementary Table S1.** Random-Forest Approximate Bayesian Computation model-choice predictions for the ACB and ASW populations. 1,000 decision trees were considered for RF prediction for the ACB and ASW respectively. Corresponding results are plotted in **Figure 3**.

| Competing<br>Model<br>Target<br>population | Afr2P-Eur2P | Afr2P-EurDE | Afr2P-EurIN | AfrDE-Eur2P | AfrDE-EurDE | AfrDE-EurIN | AfrIN-Eur2P | AfrIN-EurDE | AfrIN-EurIN |
| --- | --- | --- | --- | --- | --- | --- | --- | --- | --- |
| *ACB* | 46 | 144 | 3 | 151 | 531 | 12 | 74 | 34 | 5 |
| *ASW* | 112 | 106 | 9 | 317 | 335 | 3 | 73 | 43 | 2 |

**Supplementary Table S2.**
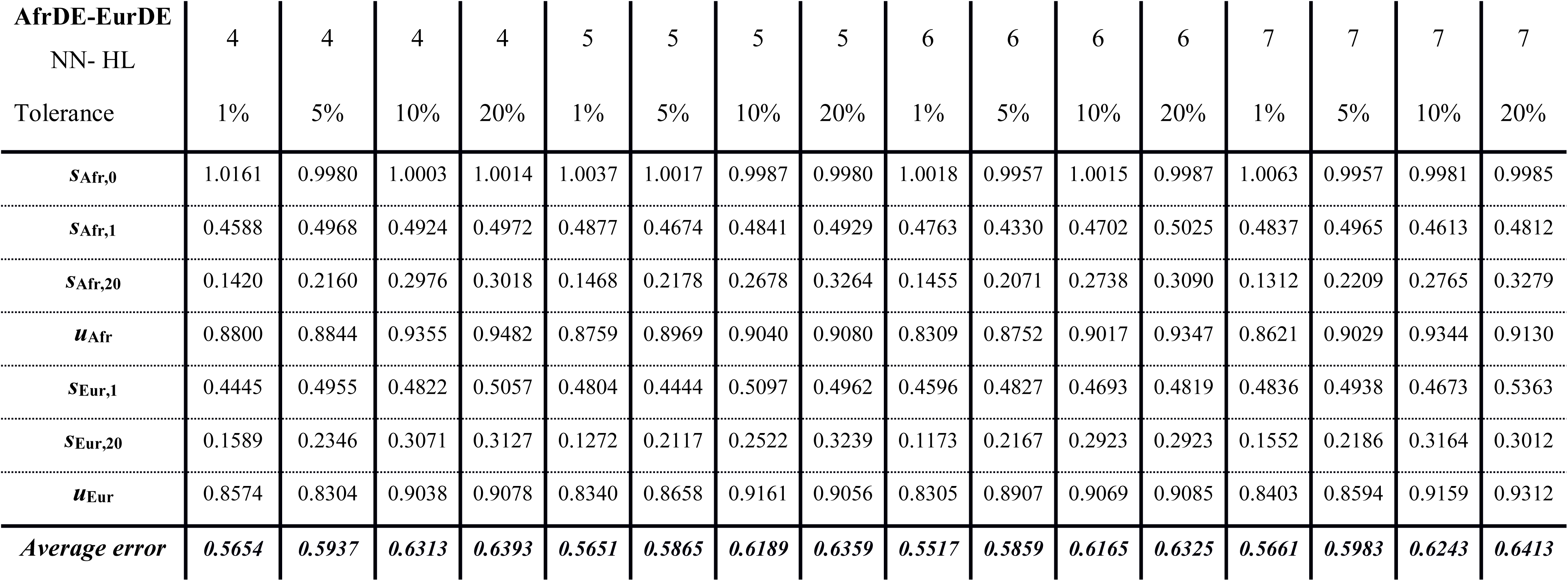
Parameter prediction cross-validation error as a function of the number of neurons in the hidden layer and the rejection tolerance rate under the AfrDE-EurDE scenario. We considered, 1,000 random simulations in turn as pseudo-observed data to estimate posterior parameter distributions, considering 4, 5, 6, or 7 neurons in the hidden layer (“NN-HL” row), and 100,000 total simulations. Tolerance levels of 0.01, 0.05, 0.1 and 0.2 were considered (“Tolerance” row). The median values of posterior parameter distributions were used as point estimates for the error calculation.

**Supplementary Table S3.** Accuracy of the 95% credibility interval estimated for posterior parameters in the vicinity of the observed ACB and ASW datasets. We provide the empirical coverage of the estimated 95% credibility interval, i.e. how many times (in percentage) the true parameter (*θ_i_*) is found inside the estimated 95% credibility interval [2.5%quantile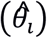; 97.5%quantile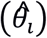], among the 1,000 posterior parameter estimations conducted using in turn the 1,000 simulations closest to our real data, separately for the ACB and ASW, as pseudo-observed datasets for four separate methods: NN estimation of the parameters taken jointly as a vector, NN estimation of the parameters taken independently, Random Forest (parameters are taken independently), and Rejection (parameters are taken independently).

| *AfrDE-EurDE*<br>*parameters* | ACB |  |  |  | ASW |  |  |  |
| --- | --- | --- | --- | --- | --- | --- | --- | --- |
|  | *NN joint* | *NN indep.* | *RF indep.* | *Rejection*<br>*indep.* | *NN joint* | *NN indep.* | *RF indep.* | *Rejection*<br>*indep.* |
| $s_{\text{Afr},0}$ | 0.956 | 0.934 | 0.929 | 0.952 | 0.952 | 0.931 | 0.937 | 0.950 |
| $s_{\text{Afr},1}$ | 0.958 | 0.929 | 0.942 | 0.968 | 0.958 | 0.914 | 0.942 | 0.963 |
| $s_{\text{Afr},20}$ | 0.964 | 0.926 | 0.956 | 0.971 | 0.963 | 0.928 | 0.960 | 0.978 |
| $u_{\text{Afr}}$ | 0.953 | 0.932 | 0.930 | 0.950 | 0.944 | 0.914 | 0.925 | 0.945 |
| $s_{\text{Eur},1}$ | 0.947 | 0.909 | 0.939 | 0.949 | 0.950 | 0.912 | 0.930 | 0.955 |
| $s_{\text{Eur},20}$ | 0.944 | 0.908 | 0.930 | 0.957 | 0.952 | 0.919 | 0.929 | 0.968 |
| $u_{\text{Eur}}$ | 0.941 | 0.919 | 0.927 | 0.943 | 0.947 | 0.928 | 0.936 | 0.952 |
| **Average**<br>**credibility**<br>**interval**<br>**accuracy** | **0.951** | **0.922** | **0.936** | **0.955** | **0.952** | **0.920** | **0.937** | **0.958** |

